# Engineering Methyl-Coenzyme M Reductase for Enhanced Methane and Carbon Dioxide Capture through Biofilm Growth

**DOI:** 10.64898/2026.06.26.734734

**Authors:** Hyeon-Ji Hwang, Mrugesh Parasa, Ruchira Mitra, Rodolfo García-Contreras, Vanesa Angarita-Zapata, Ingmar H. Riedel-Kruse, Thomas K. Wood

**Affiliations:** Department of Chemical Engineering, Pennsylvania State University, University Park, Pennsylvania, 16802-4400; Department of Microbiology and Parasitology, Faculty of Medicine, National Autonomous University of Mexico, Mexico City, 04510, Mexico; Department of Molecular and Cellular Biology, Applied Mathematics, Physics and Biomedical Engineering, University of Arizona, Tucson, Arizona, 85721

**Keywords:** ANME-1 methyl-coenzyme M reductase, protein engineering, methane capture, biofilm, reverse methanogenesis

## Abstract

Methane is a potent greenhouse gas, nearly half of which is consumed anaerobically by anaerobic methanotrophic archaea (ANME) through methyl-coenzyme M reductase (Mcr). However, ANME cannot be grown as pure cultures, and obtaining active ANME Mcr *in vitro* remains extremely challenging, preventing previous efforts to engineer this key enzyme. Here, we used directed evolution in the methanogen *Methanosarcina acetivorans* to enhance ANME-1 Mcr (Mcr_ANME-1_) activity for methane and carbon dioxide capture by selecting Mcr_ANME-1_ variants with improved growth during methane-dependent cultivation. As a result, we discovered two beneficial substitutions in the catalytic α-subunit of Mcr_ANME-1_, S60P and I154V, that increased biofilm growth as well as acetate production and methane capture. AlphaFold structural predictions suggest possible mechanistic explanations for these beneficial substitutions. These findings demonstrate that Mcr can be engineered to enhance methane and carbon dioxide capture, establishing a foundation for biological greenhouse gas mitigation and carbon utilization technologies.

## INTRODUCTION

Methane is a potent greenhouse gas, and anaerobic methanotrophic archaea (ANME) constitute one of the largest biological methane sinks on Earth, consuming up to ∼270 Tg of methane produced by methanogens [1], out of the ∼600 Tg produced annually worldwide (∼60% anthropogenic) [2]. Carbon dioxide is also a major greenhouse gas, and methane utilization by ANME can be coupled to inorganic carbon assimilation, enabling simultaneous methane and carbon dioxide capture [3]. Through anaerobic oxidation of methane (AOM), ANME play a major role in regulating global methane emissions and mitigating climate change [4]. This process also contributes to carbon cycling through the incorporation of inorganic carbon into cellular biomass and products [4].

Despite their environmental importance, ANME remain difficult to study and exploit for biotechnology because they have not been isolated as pure cultures [1,4] and grow extremely slowly in syntrophic consortia, with doubling times of approximately seven months [5]. Furthermore, methyl-coenzyme M reductase (Mcr) requires a complex maturation process involving numerous accessory proteins [6]. Consequently, mechanistic studies and protein engineering of active ANME Mcr have remained largely inaccessible [7].

Mcr from Black Sea ANME-1 (Mcr_ANME-1_) consists of a hexameric (αβγ) complex composed of α (McrA), β (McrB), and γ (McrG) subunits and catalyzes the activation of methane using the nickel porphinoid cofactor F_430_ located within McrA (**Fig. 1A**) [6,8]. During anaerobic methane oxidation, Mcr_ANME-1_ catalyzes the conversion of methane and coenzyme M/coenzyme B heterodisulfide (CoM−S−S−CoB) into methyl-coenzyme M (CH_3_−S−CoM) and coenzyme B (HS−CoB) (**Fig. 1B**) [6,8]. Assembly and activation of Mcr additionally require a large maturation system consisting of at least 17 accessory proteins [9,10]. Intriguingly, despite sharing a highly conserved overall architecture with methanogenic Mcr enzymes, Mcr_ANME-1_ performs methane activation in the opposite physiological direction of methanogens and exhibits several distinctive structural features. These differences include a modified F_430_ cofactor (methylthio-F_430_), a cysteine-rich region near the active site, and distinct patterns of post-translational amino acid modifications [11]. For example, Mcr_ANME-1_ contains thio-Glyα464, N- methyl-Hisα271, Trpα333, and Metα499 sulfoxide, whereas Mcr from *Methanosarcina* species contains thio-Gly, N-methyl-His, S-methyl-Cys, and 5-(S)-methyl-Arg [11]. These unique structural features suggest that Mcr_ANME-1_ has evolved specific adaptations for methane oxidation; however, the functional significance of these differences remains largely unknown [8].

**Fig. 1.**
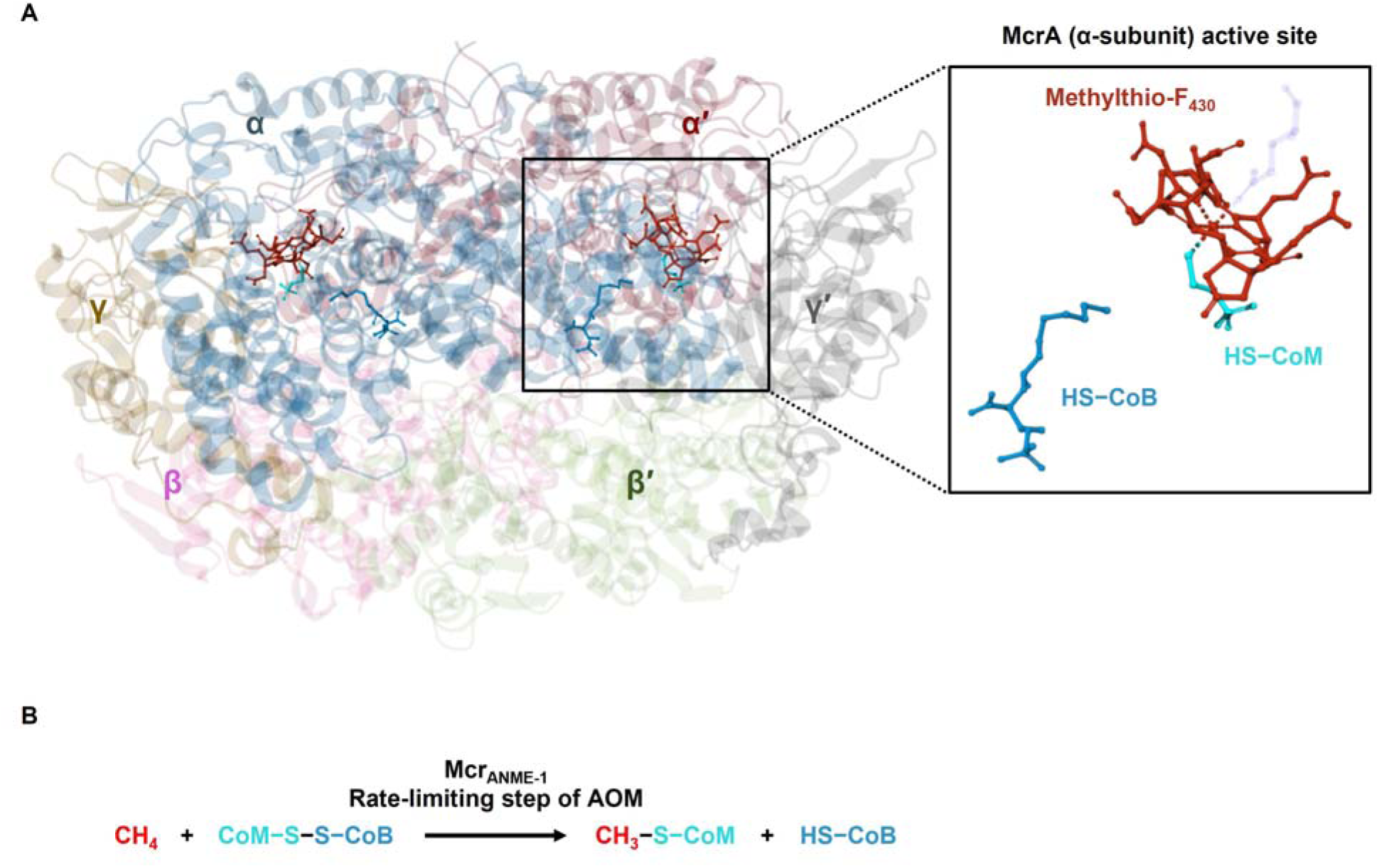
Structure and function of Mcr_ANME-1_. (**A**) Structure of Mcr_ANME-1_ showing the α (McrA), β (McrB), and γ (McrG) subunits and the active site located within the McrA subunit. Primes (′) denote the second αβγ heterotrimer in the (αβγ) Mcr_ANME-1_ complex. The catalytic cofactor methylthio-F_430_ (dark red) and the reaction-associated coenzymes HS-CoM (cyan) and HS-CoB (blue) are highlighted. Structure based on the crystal structure of Mcr_ANME-1_ (PDB ID: 3SQG). (**B**) Mcr_ANME-1_-catalyzed methane activation. Mcr_ANME-1_ catalyzes methane activation, the first committed and rate-limiting step of anaerobic methane oxidation (AOM). CH_4_, methane (red); CoM–S–S–CoB, coenzyme M/coenzyme B heterodisulfide; CH_3_–S–CoM, methyl-coenzyme M; HS–CoB, reduced coenzyme B.

To overcome the inability to cultivate ANME in pure culture and to enable biochemical studies of active ANME Mcr, *mcrBGA* from unculturable Black Sea ANME-1 was previously cloned into the methanogen *Methanosarcina acetivorans* [3]. This host provides the cofactors, maturation machinery, accessory proteins, and metabolic background required for active Mcr_ANME-1_ function because it naturally produces a homologous Mcr for methanogenesis (Mcr*_M.a._*) [12]. Importantly, heterologous expression of Mcr_ANME-1_ enabled reverse methanogenesis and supported methane-dependent growth in an anaerobic pure culture for the first time [3]. Under these conditions, methane oxidation was coupled to ferric iron reduction and incorporation of inorganic carbon into acetate according to the overall reaction:

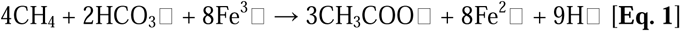

By utilizing ferric iron as the terminal electron acceptor, methane-dependent growth was demonstrated for more than three generations, establishing the first culturable anaerobic pure strain capable of growth on methane through recombinant expression of active Mcr_ANME-1_ [3]. Independent studies in *M. acetivorans* further elucidated the biochemical basis of methane-dependent metabolism coupled to ferric iron reduction and provided additional evidence supporting the reversibility of methanogenesis [13–15]. Importantly, the recombinant Mcr_ANME-1_ platform provides the first opportunity to investigate and engineer active Mcr_ANME-1_ *in vivo*.

Notably, the recombinant Mcr_ANME-1_ platform has already enabled several biotechnological applications. Acetate generated from methane capture has been converted to L-lactate through heterologous pathway engineering [16], achieving yields substantially greater than those reported for aerobic methane-utilization systems. The resulting lactic acid has applications in foods [17], cosmetics [18], pharmaceuticals [19], and biodegradable plastics [20]. The engineered strain has also been utilized to construct the first biological fuel cell that converts methane into electricity [21,22]. This methane microbial fuel cell operates with high Coulombic efficiency and achieves power and current densities of 5.2 W/m² and 7 A/m², respectively, comparable to the best microbial fuel cells utilizing conventional gaseous or non-gaseous substrates [21,22]. More recently, the platform was used to convert methane and inorganic carbon into the biofuel ethanol [23]. Collectively, these studies demonstrate the versatility of recombinant Mcr_ANME-1_ for methane-based biomanufacturing.

Despite these advances, methane activation by Mcr_ANME-1_ remains the first committed step in anaerobic methane oxidation and likely represents a major rate-limiting step in methane-dependent growth and product formation (**Fig. 1B**). Consequently, the catalytic performance of Mcr_ANME-1_ is expected to influence not only methane capture but also the efficiency of downstream methane-based biotechnological applications. Improving Mcr_ANME-1_ therefore represents a promising strategy to enhance methane utilization, carbon incorporation, and bioproduct synthesis. However, active Mcr_ANME-1_ remained inaccessible to protein engineering until the recent development of a suitable cultivation and expression platform.

Here, we report the first directed evolution of Mcr. Error-prone polymerase chain reaction (epPCR) has previously been used successfully by us to evolve enzymes including monooxygenases [24], dioxygenases [25], and epoxide hydrolases [26]. Applying this approach to *mcrA*, we generated random mutations and selected variants with enhanced methane-dependent growth, identifying beneficial substitutions in the Mcr_ANME-1_ α-subunit. Subsequent saturation mutagenesis [27] revealed S60P and I154V as improved variants that enhanced methane-dependent growth and methane utilization. These findings establish, for the first time, that active Mcr_ANME-1_ can be engineered to enhance methane utilization and methane capture. More broadly, they provide a foundation for the rational development of methane-oxidizing biocatalysts for greenhouse-gas mitigation, carbon utilization, and methane-based biomanufacturing.

## RESULTS

### Directed evolution strategy for improving Mcr_ANME-1_ function in a heterologous host

We hypothesized that Mcr_ANME-1_ activity is not fully optimized in the heterologous host *M. acetivorans* because ANME-1 evolved separately from methanogenic archaea [28], and the host Mcr*_M.a._* primarily produces methane whereas Mcr_ANME-1_ captures it [23]. Since Mcr_ANME-1_ catalyzes the first and rate-limiting step of AOM (**Fig. 1B**), improving its activity was expected to enhance methane-dependent growth in the engineered host. Under methane cultivation conditions, growth occurs predominantly as biofilms (**Fig. S1**), which facilitate electron transfer from methane oxidation to insoluble ferric iron serving as the terminal electron acceptor [3]. Therefore, biofilm formation was used as a phenotypic proxy for improved methane-dependent growth and enhanced Mcr_ANME-1_ activity. To generate functional diversity, we employed epPCR targeting *mcrA*, which encodes the catalytic α-subunit containing the active site of Mcr_ANME-1_ (**Fig. 1A**). Variants with improved methane-dependent growth were subsequently enriched through long-term cultivation under methane-selective conditions (**Fig. 2**).

**Fig. 2.**
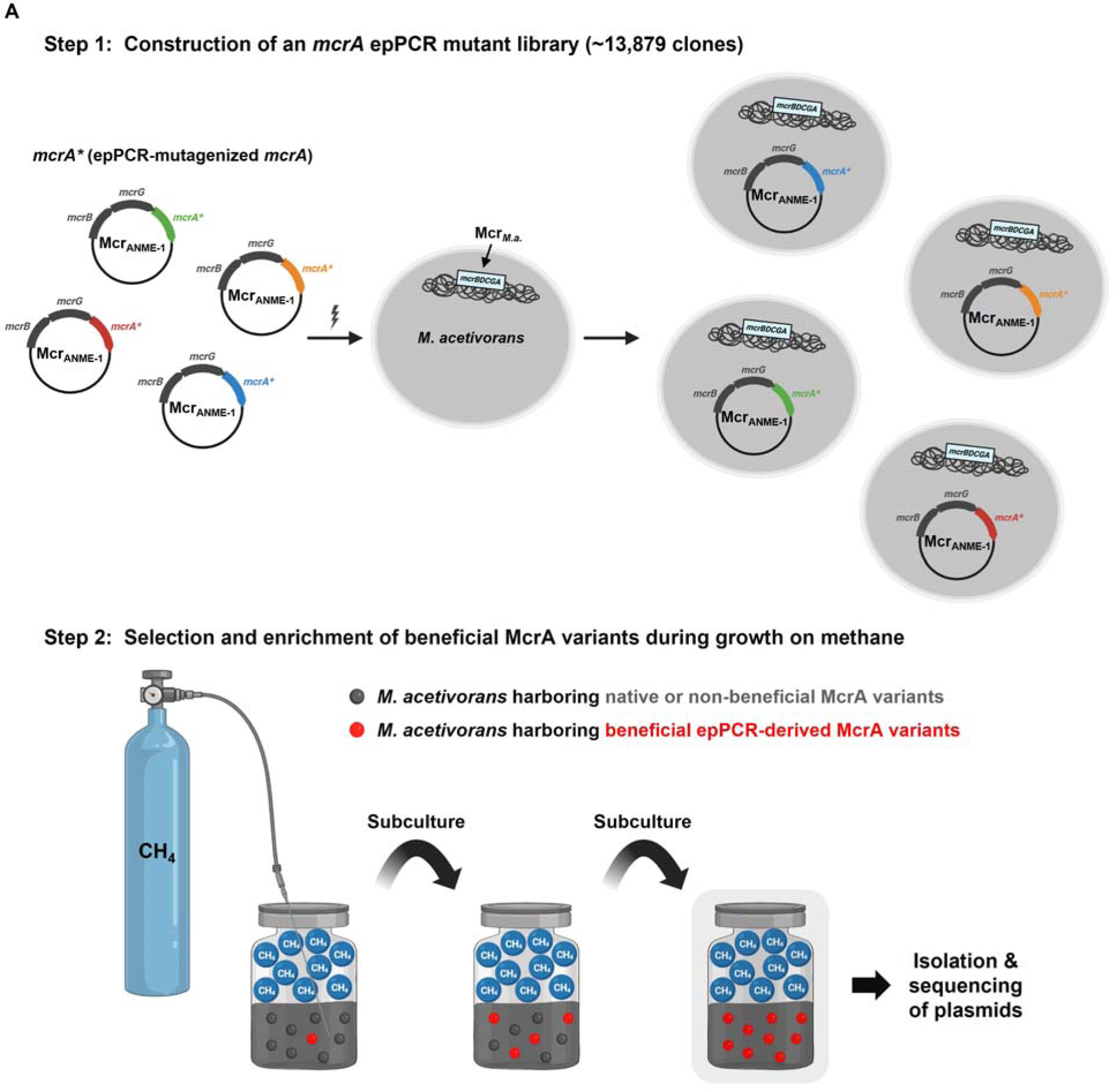
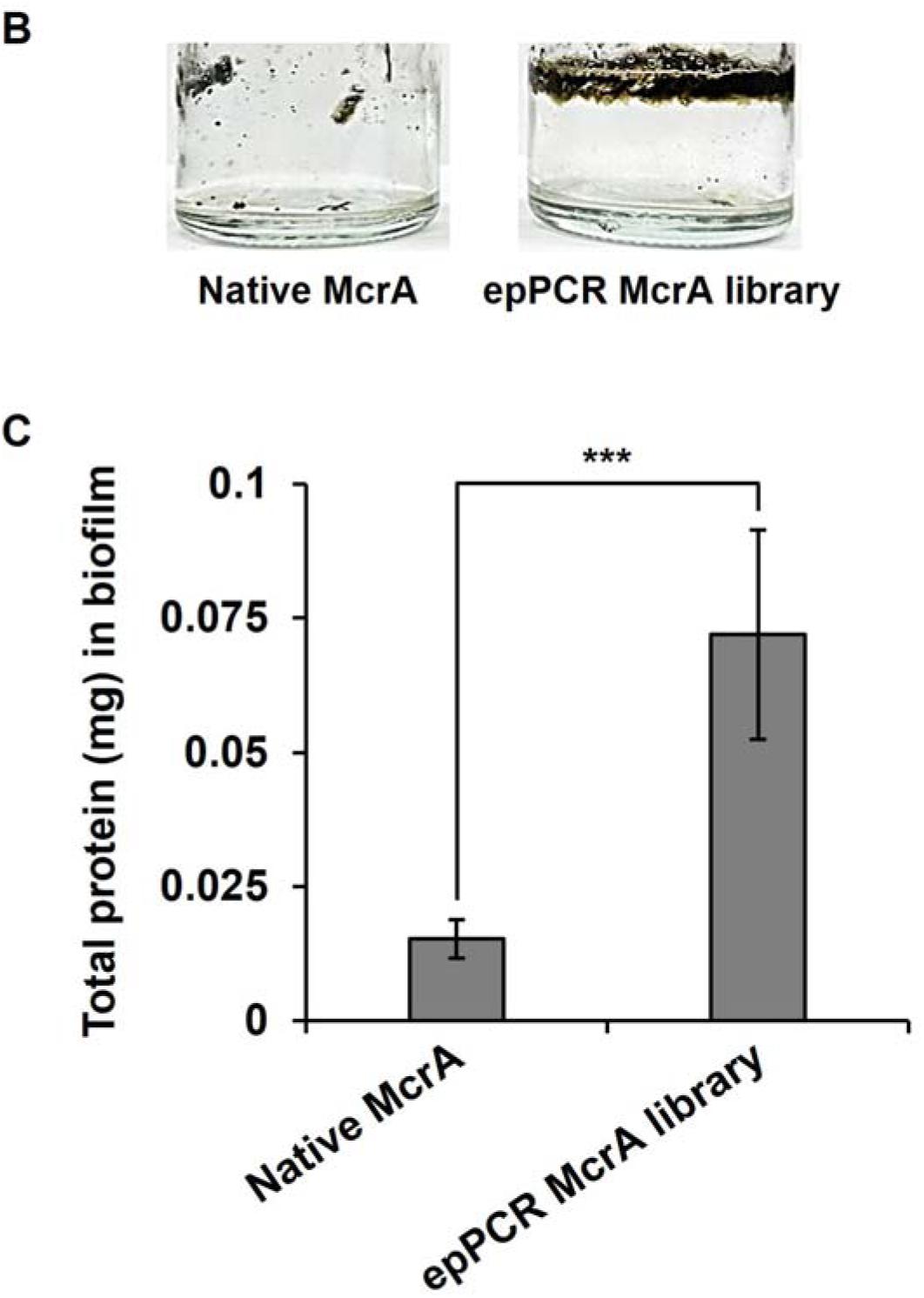
Selection of beneficial Mcr_ANME-1_ McrA variants by directed evolution and growth on methane. (**A**) Schematic of the selection and enrichment strategy used to identify beneficial Mcr_ANME-1_ McrA variants during growth on methane. An *mcrA* epPCR mutant (*) library was introduced into *M. acetivorans* by electroporation (Step 1). During serial cultivation on methane, cells carrying beneficial Mcr_ANME-1_ McrA variants became enriched relative to cells carrying native or non-beneficial variants (Step 2). Plasmids were subsequently isolated and sequenced to identify beneficial mutations. Created with BioRender.com. (**B**) Representative images showing biofilm formation at the methane–liquid interface by *M. acetivorans* producing native Mcr_ANME-1_ McrA or epPCR-derived Mcr_ANME-1_ McrA variants after 5 days of methane cultivation. (**C**) Total biofilm protein levels. Cultures were grown on methane for ∼5 days. Data are shown as mean ± standard deviation from two independent experiments. *, *P* < 0.05; ***, *P* < 0.005.

### Construction of the Mcr_ANME-1_ *mcrA* mutant library spanning mutation rates from 0.4 to 3.4%

Initial attempts to mutagenize the entire Mcr_ANME-1_ *mcrBGA* operon (∼3.9 kb) did not yield variants with improved methane-dependent growth. Therefore, we focused mutagenesis on *mcrA*, which encodes the catalytic α-subunit of Mcr_ANME-1_. To facilitate library construction, the NcoI site located in the plasmid backbone was removed, generating pES1-MAT*mcr3*-NcoI-mut (**Fig. S2**), in which *mcrA* is flanked by unique NcoI and NheI restriction sites (**Fig. S3**). epPCR conditions were optimized to target mutation frequencies ranging from approximately 0.5 to 3% by varying MgCl_2_ concentrations (1.5 to 7.5 mM), MnCl_2_ concentrations (0.09 to 0.7 mM), and dNTP ratios (dATP + dGTP : dTTP + dCTP = 1:1.15 to 1:5) (**Table S1**). Mutation frequencies were estimated by sequencing three representative clones from each mutagenesis condition. We successfully generated mutation rates spanning 0.4 to 3.4% (**Table S1**), enabling construction of libraries with varying levels of sequence diversity. After excluding colonies arising from vector self-ligation, a total of approximately 13,879 independent *mcrA* variants were obtained. The resulting library was introduced into *M. acetivorans* C2A by electroporation and subjected to methane-dependent growth selection (**Fig. 2A**).

### Long-term methane selection identifies Mcr_ANME-1_ McrA variants with 5-fold enhanced biofilm growth

To identify improved Mcr_ANME-1_ variants, the mutant library was cultivated under methane-dependent growth conditions using methane as the electron donor and ferric iron as the terminal electron acceptor. Following approximately 50 days of selection, the evolved population exhibited substantially increased biofilm formation compared with the strain expressing native Mcr_ANME-1_. Total biofilm protein increased approximately 5-fold, from 0.015 ± 0.004 mg protein for the native enzyme to 0.07 ± 0.02 mg protein for the selected population (**Fig. 2B** and **2C**). Plasmids recovered from the selected population were subsequently analyzed to identify enriched mutations. Sequence analysis of representative clones identified one clone containing only the McrA substitution S60P, as well as additional clones carrying combinations of substitutions K52R, L94P, W96R, T125I, N139D, I154V, F174S, S306T, L357Q, F360S, and L384R. Because residue I154 is located near the catalytic center of Mcr_ANME-1_, I154V was selected for further investigation together with S60P.

### Saturation mutagenesis confirms Mcr_ANME-1_ McrA S60P and I154V as major beneficial substitutions

To further evaluate the importance of residues S60 and I154 and to ensure all 19 possible substitutions were tested at these positions, saturation mutagenesis libraries were constructed for both positions. More than 300 independent clones were obtained for each library, providing approximately 99% confidence of covering all possible amino acid substitutions at each site [29]. The libraries were introduced into *M. acetivorans* C2A and subjected to methane-dependent growth selection. In parallel, *M. acetivorans* strains expressing either the S60P or I154V McrA variant were generated to directly validate the substitutions identified by directed evolution. Strains expressing S60P McrA or I154V McrA exhibited substantially increased biofilm formation relative to the native Mcr_ANME-1_ strain (0.005 ± 0.006 mg protein), reaching 0.034 ± 0.009 mg protein for S60P and 0.032 ± 0.006 mg protein for I154V, corresponding to approximately 8-fold and 7-fold increases, respectively (**Fig. 3A** and **3B**). Additionally, the S60P and I154V variants exhibited enhanced methane capture, reaching 14 ± 2 and 25 ± 1 μmol/mg, respectively, compared with 5 ± 1 μmol/mg for the native Mcr_ANME-1_ strain, representing 2.7-fold and 4.9- fold increases (**Fig. 3C**). Similarly, acetate production also increased from 1.0 ± 0.1 μmol/mg in the native Mcr_ANME-1_ strain to 2.8 ± 0.5 μmol/mg in S60P and 5.4 ± 0.4 μmol/mg in I154V, corresponding to 2.7-fold and 5.2-fold increases, respectively (**Fig. 3C**). These results confirm that both the S60P and I154V substitutions in Mcr_ANME-1_ McrA are beneficial for methane-dependent growth. Consistent with these findings, both saturation mutagenesis libraries also exhibited enhanced biofilm formation relative to the native Mcr_ANME-1_ strain, reaching 0.037 ± 0.003 mg protein for the S60 library and 0.03 ± 0.02 mg protein for the I154 library, corresponding to approximately 8-fold and 7-fold increases, respectively (**Fig. 3A** and **3B**). These results further support the importance of residues S60 and I154 in determining Mcr_ANME-1_ performance during methane-dependent growth.

**Fig. 3.**
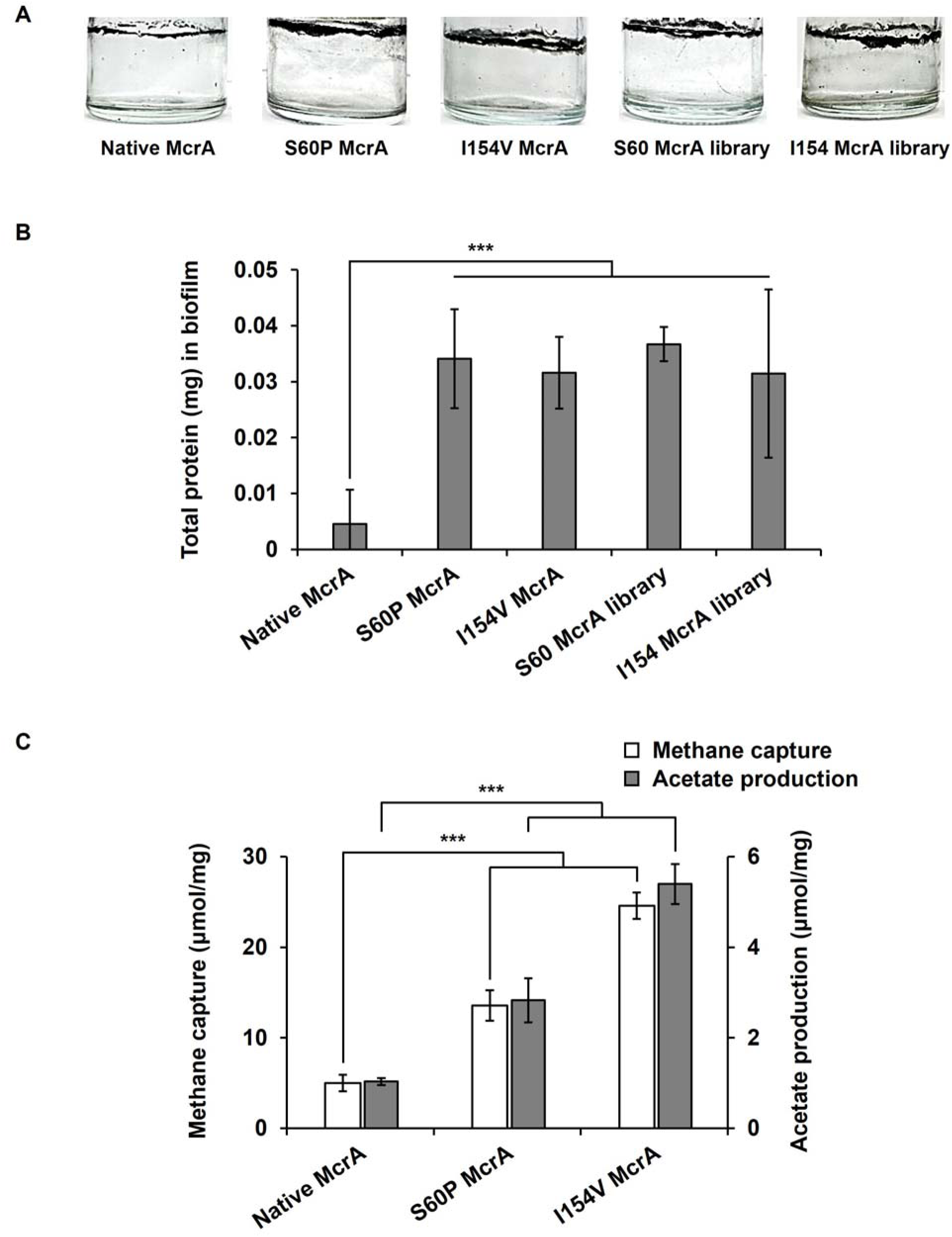
Further engineering of beneficial Mcr_ANME-1_ McrA residues by saturation mutagenesis. (**A**) Representative images of biofilm formation after ∼14 days at the methane–liquid interface by *M. acetivorans* producing native Mcr_ANME-1_ McrA, the corresponding Mcr_ANME-1_ S60P and I154V McrA variants, and the Mcr_ANME-1_ S60 and I154 McrA saturation mutagenesis libraries. (**B**) Total biofilm protein levels. Cultures were grown on methane for ∼14 days. (**C**) Methane capture (μmol/mg) and acetate production (μmol/mg) by the native Mcr_ANME-1_ strain and the Mcr_ANME-1_ S60P and I154V McrA variants. Values were normalized to cell biomass (total protein), and cultures were grown on methane for 6 days. Data are shown as mean ± standard deviation from at least three independent experiments. *, *P* < 0.05; **, *P* < 0.01; ***, *P* < 0.005.

To identify additional beneficial substitutions, plasmids recovered from the selected saturation mutagenesis populations were sequenced. No substitutions superior to S60P or I154V were identified. However, sequencing revealed a single substitution in two clones, W528C, which was not intentionally targeted during library construction. This mutation was likely introduced during PCR amplification and subsequently enriched during methane-dependent growth selection. Collectively, these results confirm that S60P and I154V are major beneficial substitutions and identify W528C as a possible additional beneficial substitution for improving Mcr_ANME-1_ performance.

### Structural modeling reveals distinct mechanisms for Mcr_ANME-1_ McrA S60P and I154V

To investigate possible mechanisms underlying improved activity, the positions of beneficial substitutions were analyzed using AlphaFold 3 structural models [30] of Mcr_ANME-1_. The I154V substitution is located approximately 9 Å from the nickel atom of the methylthio-F_430_ cofactor (**Fig. 4A**) and lies within a flexible loop region adjacent to the catalytic pocket (**Fig. S4**). Notably, residue I154 in Mcr_ANME-1_ corresponds to valine (V) in many methanogenic and methanotrophic archaeal Mcr homologs, including Mcr*_M.a._* (**Fig. 4B**). Thus, the I154V substitution represents a reversion toward the residue commonly found at the corresponding position in other archaeal Mcr enzymes. This reversion may alter the local environment of the catalytic pocket in a manner that favors methane activation. The adjacent residue Q155 is highly conserved among methanogenic and methanotrophic Mcr enzymes (**Fig. 4B**) and directly coordinates the nickel center of methylthio-F_430_ [11,31]. Previous studies have shown that this glutamine (Q) helps position the cofactor and regulates the electronic state of the nickel center [31,32]. Therefore, the close proximity of I154V to Q155 raises the possibility that the substitution influences the electronic environment of the nickel center and thereby contributes to improved catalysis. In contrast, S60P is located approximately 30.2 Å from the nickel center of methylthio-F_430_ (**Fig. 4A**). Structural modeling suggests that the substitution may alter local conformational dynamics that propagate to the catalytic region through long-range structural effects (**Fig. S4**). Sequencing of the selected saturation mutagenesis populations also identified the substitution W528C. Similar to S60P, W528C is located distal to the catalytic center, approximately 33 Å from the methylthio- F_430_ cofactor (**Fig. S5A**), yet within 5 Å of residues lining the tunnel that controls the entry of the heterodisulfide CoM–S–S–CoB and the exit of the coenzymes HS-CoM and HS-CoB (**Fig. S5B**). Replacement of the bulky aromatic tryptophan (W) residue with cysteine (C) is predicted to induce local structural rearrangements of this tunnel, thereby potentially influencing the binding and rate of transport of the cofactors into and out of the tunnel and increasing the turnover of methane.

**Fig. 4.**
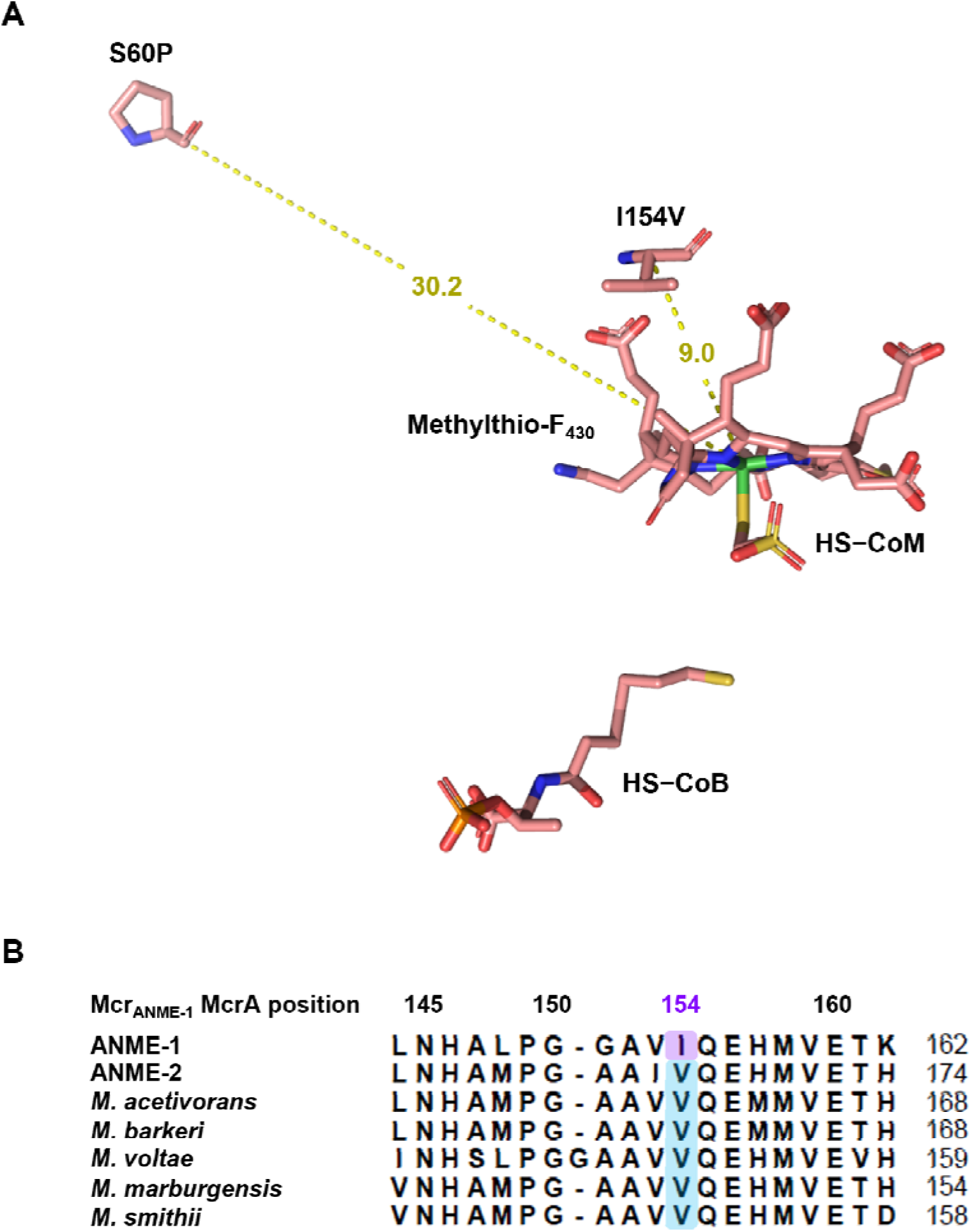
Structural context of the beneficial Mcr_ANME-1_ McrA substitutions S60P and I154V. (**A**) Distances (Å) from the nickel center of the methylthio-F_430_ cofactor to the beneficial substitutions S60P and I154V. Carbon is shown in pink, oxygen in red, nitrogen in blue, sulfur in yellow, phosphorus in orange, and nickel in green. (**B**) Comparison of the McrA amino acid sequence surrounding residue 154 among representative anaerobic methanotrophic (ANME) and methanogenic archaea. Native Mcr_ANME-1_ McrA contains an isoleucine (I; purple) at position 154, whereas a valine (V; blue) is present at the corresponding position in the other McrA homologs shown. The glutamine (Q) residue at position 155, which directly coordinates the nickel center of the methylthio-F_430_ cofactor, is conserved among all McrA homologs shown.

To further investigate whether the beneficial substitutions S60P, I154V, and W528C shared common evolutionary features, sequence comparisons were performed. Interestingly, residues corresponding to S60, I154, and W528 in Mcr_ANME-1_ McrA are occupied by smaller amino acids in the native Mcr*_M.a._* McrA sequence (A, V, and L, respectively) (**Fig. S6**). Similarly, the beneficial substitutions I154V and W528C identified in this study also reduce side-chain size relative to the native Mcr_ANME-1_ residues. This suggests that mutations selected during methane-dependent growth may partially reflect adaptation of Mcr_ANME-1_ to the intracellular environment of *M. acetivorans*. The observed shift toward host-like residues indicates that laboratory evolution favored variants that are potentially better accommodated within the heterologous host.

## DISCUSSION

To our knowledge, this study represents the first directed evolution of Mcr. Although Mcr_ANME-1_ is a highly complex enzyme that requires extensive post-translational modifications and specialized maturation machinery, we found that directed evolution successfully identified beneficial substitutions that improved methane-dependent growth in a heterologous host.

Notably, beneficial variants were recovered almost exclusively from libraries with mutation frequencies below approximately 1%, with an average mutation frequency of approximately 0.6%. This observation suggests that Mcr_ANME-1_ is highly sensitive to excessive mutational burden and that higher-error variants likely suffered loss of function due to disruption of protein folding, maturation, or catalysis. The enrichment of low-error variants highlights the importance of balancing diversity and functionality during engineering of complex archaeal enzymes.

An additional observation was that growth occurred predominantly as biofilms located at the methane–liquid interface (**Fig. 2B**). Because methane has limited solubility in aqueous media, localization at the gas–liquid interface likely enhances methane accessibility and supports more efficient methane utilization. The increased biofilm formation observed in strains expressing engineered Mcr_ANME-1_ variants therefore likely reflects improved methane-dependent growth rather than simply altered surface attachment.

Two Mcr_ANME-1_ McrA substitutions, S60P and I154V, consistently enhanced biofilm formation and were independently validated through saturation mutagenesis and reconstruction experiments. Particularly notable was the I154V substitution, which reverts an ANME-specific isoleucine (I) residue to the valine (V) present in many methanogenic and methanotrophic archaeal Mcr homologs. Given its proximity to the methylthio-F_430_ catalytic center and the conserved neighboring Q155 residue, this substitution may influence the local structural environment surrounding the active site and thereby improve catalytic efficiency. Q155 directly coordinates the nickel center of methylthio-F_430_ and has been proposed to regulate its electronic state [31]. Therefore, the close proximity of I154V to Q155 raises the possibility that this substitution influences the electronic environment of the catalytic center and thereby enhances methane activation.

A striking observation was that the I154V and W528C substitutions reduce side-chain size relative to the native Mcr_ANME-1_ residues, similar to the corresponding residues in native Mcr*_M.a._*. These observations suggest that selection during directed evolution may have partially optimized Mcr_ANME-1_ for function within the intracellular environment of the heterologous host. Thus, in addition to improving methane-dependent growth, the selected mutations may reflect adaptive interactions between Mcr_ANME-1_ and the cellular environment of *M. acetivorans*.

Future quantum chemical calculations comparing transition-state energies with either I or V at position 154 could help quantify the energetic consequences of the I154V substitution and provide further mechanistic insight into methane activation. However, such analyses are currently limited by the lack of an experimentally resolved structure of heterologously expressed Mcr_ANME-1_ in *M. acetivorans*. Structural determination of the engineered enzyme will therefore be an important objective for future studies.

An additional consideration related to the beneficial substitutions is that unlike in methanogens, where methanogenesis is initiated by the entry of the cofactors, methyl-coenzyme M and coenzyme B [33], in methane-consuming archaea, the heterodisulfide CoM–S–S–CoB enters the tunnel first [34]. This change in cofactor directionality likely imposes distinct structural constraints on residues lining the transport tunnel (**Fig. S6**). In our case, without the substitutions, a non-optimal tunnel within Mcr_ANME-1_ might be present when expressed in *M. acetivorans* due to (i) incorporation of the host-derived F_430_ cofactor instead of the native Mcr_ANME-1_ methylthio-F_430_ cofactor, (ii) the possible lack of certain post-translational modifications in *M. acetivorans* that are native to ANME-1, or (iii) the possible incorporation of novel post-translational modifications native to *M. acetivorans*. Furthermore, recent experiments have shown that hybrid Mcr structures can form between subunits of the host Mcr*_M.a._*and the heterologously expressed Mcr [35]. The W528C substitution might adjust tunnel dynamics to compensate for these effects when expressed in *M. acetivorans*. The S60P and I154V substitutions, though not located near the tunnel, might also reorganize tunnel architecture through long-range effects. To quantify the impact of the substitutions identified in this study on the rate of transport of cofactors into and out of the tunnel, molecular dynamics simulations and NMR studies are needed.

In our study, *M. acetivorans* was grown at its optimal growth temperature of 39 °C and at atmospheric pressure (1 bar), whereas native Mcr_ANME-1_ originates from deep-sea sediments where temperatures are near freezing and pressures are two to three orders of magnitude higher than those at sea level [36]. Proteins from psychrophilic and piezophilic organisms are generally more flexible than those from their mesophilic counterparts [37–39]. When proteins from psychrophilic or piezophilic organisms are transferred to mesophilic hosts, compensatory adaptations may be required to offset the increased molecular fluctuations associated with higher temperatures. ANME-derived Mcr proteins were expressed at higher levels at room temperature than at 37 °C when heterologously produced in model methanogens, suggesting improper folding and subsequent degradation at higher temperatures [35]. Both the S60P and W528C substitutions may increase protein rigidity relative to the native Mcr_ANME-1_. While the S60P substitution introduces proline, which has exceptionally high rigidity [40], the W528C substitution introduces a cysteine residue that may reduce local flexibility and could potentially participate in disulfide bond formation [41]. Although no cysteine residue is located within disulfide-bonding distance (∼2 Å) of W528C in the Mcr_ANME-1_ McrA structure, the intriguing possibility of chimeric Mcr proteins forming with subunits from *M. acetivorans* [35] does not rule out disulfide bond formation.

Beyond identifying beneficial mutations, this work also establishes a general framework for engineering enzymes that catalyze anaerobic methane activation. The ability to evolve Mcr_ANME-1_ *in vivo* creates opportunities to improve methane capture and enhance methane conversion into valuable products including acetate, ethanol, lactate, hydrogen, and electricity. More broadly, these results transform Mcr_ANME-1_ from a difficult-to-study environmental enzyme into an engineerable biocatalyst and provide one of the first experimental links between Mcr_ANME-1_ sequence variation and methane-dependent growth phenotypes.

Importantly, this work was only possible because active Mcr_ANME-1_ can be functionally expressed in *M. acetivorans*, whereas the original ANME organisms remain unavailable as pure cultures. Consequently, the platform described here not only enables engineering of Mcr_ANME-1_ but also provides a unique system for experimentally investigating the structure–function relationships of methane-oxidizing Mcr enzymes. As such, this study establishes a foundation for future efforts in archaeal protein engineering and methane biotechnology.

## CONCLUDING REMARKS

Methyl-coenzyme M reductase (Mcr) is the key enzyme responsible for biological methane activation and production, yet its engineering has remained largely unexplored. The structural complexity of Mcr, together with its unique post-translational modifications, specialized maturation machinery, and requirement for the nickel-containing cofactor F_430_, has limited efforts to directly modify and optimize its catalytic properties. Consequently, most previous studies have focused on Mcr assembly, activation, maturation, or post-translational modification rather than engineering of enzyme function itself. Here, we report the first directed evolution of Mcr to improve its activity. By selecting for enhanced Mcr_ANME-1_ activity through growth on methane, we discovered two important amino acid substitutions that increased Mcr_ANME-1_ activity by up to 8-fold in a heterologous host. These findings establish that Mcr is amenable to laboratory evolution and demonstrate that methane-activating enzymes can be experimentally optimized. As methane is both a potent greenhouse gas and an abundant carbon feedstock, the ability to engineer Mcr opens new opportunities for biological methane capture and methane valorization. More broadly, this work provides a foundation for future reverse methanogenesis-based biotechnologies. Key opportunities, challenges, and unresolved questions for the future development of engineered Mcr are summarized in the Outstanding Questions section.

## TECHNOLOGY READINESS

Heterologous production of Mcr_ANME-1_ has already been used to convert methane into acetate, lactate, ethanol, and electricity, with patents already established for the electricity-generating biofuel cell application. The evolved Mcr_ANME-1_ variants described here represent an early proof-of-concept for engineering methane-activating enzymes to improve biological methane capture and conversion. These enhancements in Mcr_ANME-1_ activity should increase the competitiveness of methane valorization technologies; however, substantially larger improvements in Mcr activity, methane uptake rates, and overall process efficiency will be required before these applications become economically viable at industrial scale. Nevertheless, this study establishes a pathway toward improved biological methane capture, greenhouse gas mitigation, and methane-to-chemicals technologies.

## OUTSTANDING QUESTIONS

- What molecular mechanisms underlie the beneficial effects of the substitutions selected here for growth on methane via heterologous production of Mcr_ANME-1_?
- Can methane capture be further increased by combining evolved Mcr_ANME-1_ with our recently developed RNA inhibition technology that enhances methane uptake and conversion to acetate and ethanol by silencing host Mcr*_M.a._*, which primarily produces methane and likely competes with Mcr_ANME-1_?
- What role do differences in the nickel porphinoid cofactor F_430_ between the methanogenic host and ANME-1 play in controlling methane capture and methane generation?
- Can the addition of mediators further increase methane capture by the evolved Mcr_ANME-1_?
- Will combining the beneficial substitutions further enhance Mcr_ANME-1_ activity?

## RESOURCE AVAILABILITY

### Lead contact

Requests for further information and resources should be directed to and will be fulfilled by the lead contact, Prof. Thomas K. Wood.

### Materials availability

All unique strains, plasmids, and materials generated in this study are available from the lead contact upon reasonable request. Distribution of these materials for non-commercial research purposes may require a materials transfer agreement (MTA). Commercial use of these materials may be subject to additional licensing requirements.

### Data and code availability

All data supporting the findings of this work are available within the article and its supplemental information files. This study did not generate any standardized dataset requiring deposition in a public repository. This study does not report original code. Any additional information required to reanalyze the data reported in this paper is available from the lead contact upon reasonable request.

## AUTHOR CONTRIBUTIONS

**Hwang, Hyeon-Ji:** Investigation (Lead), Methodology (Lead), Formal analysis (Lead), Writing - original draft (Equal), Writing - review & editing (Equal)

**Mrugesh Parasa:** Formal analysis – Structural analysis (Lead), Visualization (Lead), Writing – original draft (supporting), Writing - review & editing (supporting)

**Ruchira Mitra:** Investigation – GC analysis (Lead), Formal analysis (Supporting)

**Garcia-Contreras, Rodolfo:** Investigation (Supporting)

**Vanesa Angarita-Zapata:** Investigation (Supporting)

**Ingmar H. Riedel-Kruse:** Funding acquisition (Lead), Project administration (Equal), Supervision (Equal), Writing - review & editing (Supporting)

**Wood, Thomas K.** *(Corresponding Author)*: Conceptualization (Lead), Formal analysis (Equal), Funding acquisition (Equal), Project administration (Lead), Supervision (Lead), Writing - original draft (Lead), Writing - review & editing (Lead)

## ACKNOWLEDGMENTS

This work was supported by National Science Foundation grant 2229070 (NSF-FMRG). In addition, RGC was supported by a PASPA-DGAPA-UNAM grant for sabbatical stays and by a PAPIIT-UNAM grant IN200224. We thank Jine Li for constructing pES1-MAT*mcr3*-NcoI-mut. We also thank J. Cuello, R. Sierra Alvarez, and members of the Riedel-Kruse Lab for stimulating discussions.

## METHOD DETAILS

### Strains, culture conditions, and plasmids

The strains and plasmids used in this study are listed in **Table S2**. *M. acetivorans* C2A and its engineered strains were used in most experiments. For plasmid propagation, purification, and library construction, *Escherichia coli* DH5α λ*pir* (a host for replication of plasmids containing the R6K origin) was used. For methanol pre-cultures, all *M. acetivorans* strains were grown anaerobically in high-salt (HS) medium [3] supplemented with 2.5 g/L yeast extract (hereafter referred to as HSYE), 50 mM 1,4-piperazinediethanesulfonic acid (PIPES), and 0.5% (vol/vol) methanol as the primary carbon and energy source at pH 6.5 and 39 °C with shaking at 200 rpm until maximal growth was reached (∼7 days; optical density at 600 nm (OD_600_) ≈ 1 for the wild-type C2A strain and OD_600_ ≈ 0.2 for engineered strains). For growth on methane, fully-grown methanol cultures were harvested by centrifugation at 8,000 rpm for 15 min, washed twice with the original culture volume of HSYE medium supplemented with 50 mM PIPES to remove residual methanol, and inoculated into 12 to 15 mL of HSYE medium supplemented with 50 mM PIPES, 10 mM FeCl_3_, and 100 mg/L humic acid. For high-cell-density methane cultures (OD_600_ ≈ 20) used for gas chromatography (GC) analysis, HS medium supplemented with 50 mM PIPES and 10 mM FeCl_3_ (without yeast extract or humic acid) was used to minimize potential effects of these components on methane consumption and product formation. Methane gas (99%) was sparged through the culture medium for 5 min at 20 psi and a flow rate of ∼226 mL/min through an inlet needle, while an outlet needle was used to maintain atmospheric pressure and replace the headspace gas with methane. Cultures were then incubated in an inverted position at 39 °C with shaking at 250 rpm to minimize gas leakage. *E. coli* DH5α λ*pir* was grown in Luria–Bertani (LB; tryptone 10 g/L, yeast extract 5 g/L, and NaCl 10 g/L) medium at 37 °C with vigorous shaking at 250 rpm. For solid media, agar was added to a final concentration of 1.5% (wt/vol). Cell growth was monitored by measuring OD_600_. Antibiotics were used at the following concentrations for plasmid maintenance: puromycin (2 μg/mL) for *M. acetivorans* and ampicillin (100 μg/mL) for *E. coli* [3].

### Construction of pES1-MAT*mcr3*-NcoI-mut

In pES1-MAT*mcr3*, one NcoI restriction site is located upstream of *mcrA*, whereas a second NcoI site is present in the plasmid backbone (**Fig. S2**). To facilitate cloning of epPCR-mutagenized *mcrA* using the NcoI and NheI restriction sites, the backbone NcoI site was disrupted (**Fig. S2**). Briefly, pES1-MAT*mcr3* was digested with HpaI and self-ligated to generate a subclone containing the backbone NcoI site. The resulting plasmid was digested with NcoI, and the 5′ overhangs were filled in using T4 DNA polymerase, resulting in insertion of four nucleotides (CATG) and disruption of the NcoI recognition sequence. The mutated fragment was then cloned into the XbaI and KpnI sites of pES1-MAT*mcr3* to generate pES1-MAT*mcr3*-NcoI-mut (**Fig. S3**). Complete plasmid sequencing by Plasmidsaurus was used to confirm the mutation.

### Construction of the Mcr_ANME-1_ *mcrA* epPCR library

To generate genetic diversity throughout the *mcrA* gene for directed evolution, epPCR was performed using a modified version of a previously described protocol [42]. The Mcr_ANME-1_ *mcrA* gene encoding the catalytic α-subunit (McrA) was amplified from pES1-MAT*mcr3*-NcoI-mut. epPCR was conducted under 10 different conditions by varying MgCl_2_ (1.5 to 7.5 mM) and MnCl_2_ (0.09 to 0.7 mM) concentrations (**Table S1**). The target error rate was approximately 0.5 to 3%, sufficient to introduce functional diversity without severely compromising protein folding or activity. MgCl_2_ and MnCl_2_ concentrations were adjusted to modulate DNA polymerase fidelity, and an imbalanced dNTP ratio (dATP + dGTP : dTTP + dCTP = 1:1.15 to 1:5) was used to promote nucleotide misincorporation. Primers (25 pmol each; **Table S3**) and 10 ng of template DNA were used in each PCR reaction. The resulting PCR products were cloned into the pES1-MAT*mcr3*-NcoI-mut shuttle vector using the NcoI and NheI restriction sites (**Fig. S3**). For each epPCR condition, 3 clones containing inserts of the expected size were randomly selected and sequenced by Plasmidsaurus to estimate mutation rates. The remaining colonies from all epPCR conditions (∼13,879 clones, excluding colonies arising from vector self-ligation) were scraped, pooled, and cultured to an OD_600_ of ∼1.0. Plasmid DNA was then isolated from the pooled culture, and the resulting *mcrA* epPCR plasmid library was introduced into *M. acetivorans* C2A by electroporation (**Fig. 2A**).

### Saturation mutagenesis

To optimize enzyme activity and identify the most active McrA variants, saturation mutagenesis was performed using a modified version of a previously described protocol [29]. Positions S60 and I154, identified as beneficial hotspots by epPCR, were individually subjected to saturation mutagenesis using primers containing NNS codons (N = A, T, G, or C; S = G or C) (**Table S3**). To improve sequence-binding specificity, each degenerate primer contained at least 15 bp of non-degenerate sequence flanking the NNS codon, resulting in a total primer length of approximately 33 bp (**Table S3**). To improve PCR efficiency and facilitate library construction, a two-step PCR strategy was employed (**Fig. S7**) using the primers listed in **Table S3** rather than direct amplification of the entire ∼14.3 kb pES1-MAT*mcr3*-NcoI-mut plasmid. In the first round of PCR, two overlapping fragments were amplified separately: 482 bp and 1,752 bp for S60 saturation mutagenesis (**Fig. S7A**), or 764 bp and 1,470 bp for I154 saturation mutagenesis (**Fig. S7B**). The two fragments shared a 33-mer overlap containing the NNS codon used for saturation mutagenesis. In the second round of PCR, the overlapping fragments were assembled and amplified using the nested primers IF-mcrAup and IR-mcrAdown, generating the final 2,007 bp mutagenized product. The resulting PCR product was subsequently cloned into pES1-MAT*mcr3*-NcoI-mut using the NcoI and NheI restriction sites (**Fig. S3**). For each saturation mutagenesis library, 4 clones containing inserts of the expected size were randomly selected and sequenced by Plasmidsaurus to confirm successful mutagenesis and assess codon diversity within the library. Because screening 292 colonies has been reported to provide 99% confidence for coverage of all possible outcomes from single-site random mutagenesis [29], more than 300 clones from each saturation mutagenesis library were used for electroporation into *M. acetivorans* C2A. As described for the epPCR libraries, the remaining colonies from each saturation mutagenesis library (>300 clones, excluding colonies arising from vector self-ligation) were scraped, pooled, and cultured to an OD_600_ of ∼1.0. Plasmid DNA was then isolated from the pooled cultures, and the resulting S60 and I154 saturation mutagenesis libraries were introduced into *M. acetivorans* C2A by electroporation.

### Generation of engineered *M. acetivorans* strains

All engineered *M. acetivorans* strains were constructed by electroporation of plasmids under anaerobic conditions. Briefly, *M. acetivorans* C2A was grown anaerobically in HSYE medium (pH 6.5) supplemented with 50 mM PIPES and 0.5% (vol/vol) methanol at 39 °C with shaking at 200 rpm until maximal growth was reached (∼7 days; OD_600_ ≈ 1). Approximately 10 mL of culture was harvested by centrifugation at 8,000 rpm for 15 min, washed twice with the same volume of 0.85 M sucrose at 0 °C, and resuspended in 1/100 of the original culture volume of 0.85 M sucrose at 0 °C. Electroporation was carried out using a Gene Pulser and Pulse Controller (Bio-Rad, Hercules, CA, USA). Cells were mixed with 5 μg of plasmid DNA, and the mixtures were transferred into 0.1-cm electroporation cuvettes and incubated on ice for 1 h prior to pulse application (capacitance, 25 μF; voltage, 1.25 kV; resistance, 200 Ω) under anaerobic conditions. Immediately after electroporation, 5 mL of HSYE medium (pH 6.5) supplemented with 50 mM PIPES and 0.5% (vol/vol) methanol was added, and the cell suspension was transferred to a vial and incubated at 39 °C for 24 h prior to transfer to selective liquid medium (HSYE medium supplemented with 50 mM PIPES, 0.5% [vol/vol] methanol, and 2 μg/mL puromycin). *M. acetivorans* transformation was confirmed by either plasmid DNA isolation or water lysis heat treatment (90 °C for 60 min), followed by PCR amplification of *mcrA* using the primers listed in **Table S3**.

### Selection and enrichment during growth on methane

To select and enrich beneficial Mcr_ANME-1_ McrA variants, methane cultivation was initiated using equal amounts of cells (OD_600_ ≈ 0.1 to 0.2 or total protein ≈ 0.2 mg) to ensure comparable starting populations. Strains were cultivated in medium containing methane, carbonate, and Fe^3+^, and beneficial variants were enriched based on improved growth during methane-dependent cultivation (**Fig. 2A**). For the epPCR libraries, repeated rounds of methane cultivation were performed for up to 50 days. For the saturation mutagenesis libraries, selection was conducted for up to 3 weeks under the same methane growth conditions. Mcr activity was evaluated indirectly by monitoring biofilm formation, and biofilm biomass was quantified by measuring total biofilm protein using the Bradford assay, as described below. To minimize the effects of nutrient depletion and enable quantitative comparison among strains, biofilm protein levels were determined after short-term cultivation (5 to 14 days) using equal amounts of inoculated cells.

### Biofilm quantification and total biomass determination in methane cultures

Direct OD_600_ measurements were not used to assess growth because black FeS precipitates formed through reactions between Fe^3+^ (from FeCl_3_) and sulfide (from Na_2_S·9H_2_O), preventing accurate OD measurements. Therefore, biofilm formation was used as an indirect proxy for Mcr activity. For biofilm quantification, biofilms were scraped from vial walls, resuspended in 500 μL distilled water (DW), and lysed by sonication on ice for 4 min using alternating cycles of 30 s sonication and 30 s rest. Total biofilm protein was quantified by mixing 10 μL of lysate with 200 μL of Quick Start Bradford 1× Dye Reagent (Bio-Rad), followed by incubation at room temperature for 20 min and measurement of absorbance at 595 nm. Protein concentrations were determined using a bovine serum albumin (BSA) standard curve. Samples were diluted and reanalyzed when necessary to ensure that absorbance values remained within the linear range of the assay.

For high-cell-density methane cultures (OD_600_ ≈ 20) used for GC analysis, cell growth was assessed by quantification of total biomass protein as described previously [23]. Briefly, 100 μL of methane culture was centrifuged at 6,000 × g for 5 min. Culture supernatants were retained, while cell pellets were resuspended in an equal volume of DW, vortexed for 30 s, and heated at 90 °C for 60 min to lyse cells. Protein concentrations in both culture supernatants and cell lysates were quantified using the Bradford assay described above and summed to determine total biomass protein. The resulting total protein values were used to normalize methane consumption and acetate production.

### Protein structure modeling

Structural models of Mcr_ANME-1_ McrA variants were generated using AlphaFold 3. Predicted structures were superimposed onto the holoenzyme crystal structure of Mcr_ANME-1_ purified from Black Sea microbial mats (PDB ID: 3SQG) using PyMOL (Schrödinger, LLC). Distances between selected atoms, including distances to the nickel center of the methylthio-F_430_ cofactor, were measured using the Measurement Wizard in PyMOL. Molecular graphics and structural visualizations were generated using PyMOL. Caver Web 2.0 was used for identifying tunnels in MCR_ANME-1_ [43].

### Amino acid sequence alignments

Amino acid sequences of McrA homologs were retrieved from the UniProt and GenBank databases. Sequences included *M. acetivorans* (Q8THH1), *M. barkeri* (P07962), *Methanococcus voltae* (P11559), *Methanothermobacter marburgensis* (P11558), *Methanobrevibacter smithii* (D2ZPL4), ANME-1 (FP565147.1), and ANME-2 (MDF1530863.1). Multiple sequence alignments were generated using the ClustalW algorithm implemented in BioEdit. The resulting alignments were used to compare residue conservation among methanogenic and anaerobic methanotrophic archaeal McrA homologs. The complete sequence alignment is provided in **Fig. S8**.

### GC analysis of methane and acetate

Engineered *M. acetivorans* C2A strains harboring pES1-MAT*mcr3*-NcoI-mut-S60P (hereafter S60P) or pES1-MAT*mcr3*-NcoI-mut-I154V (hereafter I154V) were cultivated under methane growth conditions as described above. Briefly, fully grown methanol cultures were harvested, concentrated, and washed prior to inoculation into methane cultures, followed by incubation for 6 days. Prior to GC analysis, portions of the cultures were harvested for total protein measurements. All GC analyses were performed using an Agilent 6890N gas chromatograph with nitrogen as the carrier gas, as described previously [23]. Methane was identified according to its retention time, and concentrations were determined by comparison with standards. Acetate concentrations were determined by acidifying culture supernatants with 1% (vol/vol) formic acid, and the acidified supernatant was injected. Calibration curves were prepared with sodium acetate standards using the same medium composition as in the experiments. All GC sample injections were performed manually in triplicate using a high-precision syringe (Hamilton Co., Reno, NV, USA) to ensure reproducible peak areas.

## QUANTIFICATION AND STATISTICAL ANALYSIS

All experiments were performed with at least two independent cultures unless otherwise noted. Statistical significance was determined using a Student’s *t*-test assuming equal variances in Microsoft Excel. *P* < 0.05 was considered statistically significant. Data are presented as mean ± standard deviation. Exact *p*-values are provided in the figure legends.

## Supporting Information

## Supporting Information

**Table S1.** epPCR conditions used for construction of the Mcr_ANME-1_ *mcrA* mutant libraries and the corresponding mutation rates. Mutation rates were estimated from sequencing analysis of randomly selected clones from each epPCR condition. Error rates were calculated based on nucleotide substitutions identified in sequenced *mcrA* amplicons. A total of 13,879 epPCR-derived *mcrA* clones were introduced into *M. acetivorans* C2A by electroporation and screened for improved activity. *, colonies arising from vector self-ligation were excluded.

| epPCR conditions | 1 | 2 | 3 | 4 | 5 | 6 | 7 | 8 | 9 | 10 |
| --- | --- | --- | --- | --- | --- | --- | --- | --- | --- | --- |
| MgCl<sub>2</sub> (mM) | 5 | 5 | 7.5 | 7.5 | 5 | 5 | 5 | 5 | 5 | 1.5 |
| MnCl<sub>2</sub> (mM) | 0.35 | 0.5 | 0.35 | 0.7 | 0.35 | 0.35 | 0.2 | 0.13 | 0.09 | 0.35 |
| dATP + dGTP : dTTP + dCTP | 1:5 | 1:5 | 1:5 | 1:5 | 1:2 | 1:1.5 | 1:5 | 1:5 | 1:5 | 1:5 |
| Error rate (%) | 1.8 | 2.5 | 2.7 | 3.4 | 1.3 | 1 | 0.9 | 0.8 | 0.7 | 0.4 |
| Mutation rate (mutations/kb) | 18 | 25 | 27 | 33 | 13 | 10 | 9 | 9 | 8 | 4 |
| Total number of *mcrA* epPCR clones screened | 13,879* |  |  |  |  |  |  |  |  |  |

**Table S2.** Strains and plasmids used in this study. Amp^R^, ampicillin resistance; Puro^R^, puromycin resistance.

| Name | Genotype | Source |
| --- | --- | --- |
| ***Methanosarcina acetivorans*** |  |  |
| C2A | Wild-type strain of *M. acetivorans* | DSMZ<br>(DSM 2834) |
| ***Escherichia coli*** |  |  |
| DH5α λpir | F<sup>-</sup> endA1 glnV44 thi-1 recA1 relA1 gyrA96 deoR nupG<br>ϕ80dlacZΔM15 Δ(*lacZYA-argF*)U169 hsdR17(r<sub>K</sub><sup>-</sup> m<sub>K</sub><sup>+</sup>) λpir | Lab<br>collection |
| **Plasmids** |  |  |
| pES1-MATmcr3 | Amp<sup>R</sup>, Puro<sup>R</sup>, R6K ori, C2A ori, P<sub>mcr_ANME-1</sub>::*mcrBGA*<sub>ANME-1</sub> | [3] |
| pES1-MATmcr3-NcoI-mut | Amp<sup>R</sup>, Puro<sup>R</sup>, R6K ori, C2A ori, P<sub>mcr_ANME-1</sub>::*mcrBGA*<sub>ANME-1</sub>, backbone NcoI site removed | This study |
| pES1-MATmcr3-NcoI-mut-epmcrA | pES1-MATmcr3-NcoI-mut carrying an error-prone PCR-derived *mcrA* library | This study |
| pES1-MATmcr3-NcoI-mut-S60P | pES1-MATmcr3-NcoI-mut carrying McrA S60P | This study |
| pES1-MATmcr3-NcoI-mut-I154V | pES1-MATmcr3-NcoI-mut carrying McrA I154V | This study |
| pES1-MATmcr3-NcoI-mut-W528C | pES1-MATmcr3-NcoI-mut carrying McrA W528C | This study |
| pES1-MATmcr3-NcoI-mut-S60 library | pES1-MATmcr3-NcoI-mut carrying an McrA S60 saturation mutagenesis library | This study |
| pES1-MATmcr3-NcoI-mut-I154 library | pES1-MATmcr3-NcoI-mut carrying an McrA I154 saturation mutagenesis library | This study |

**Table S3.**
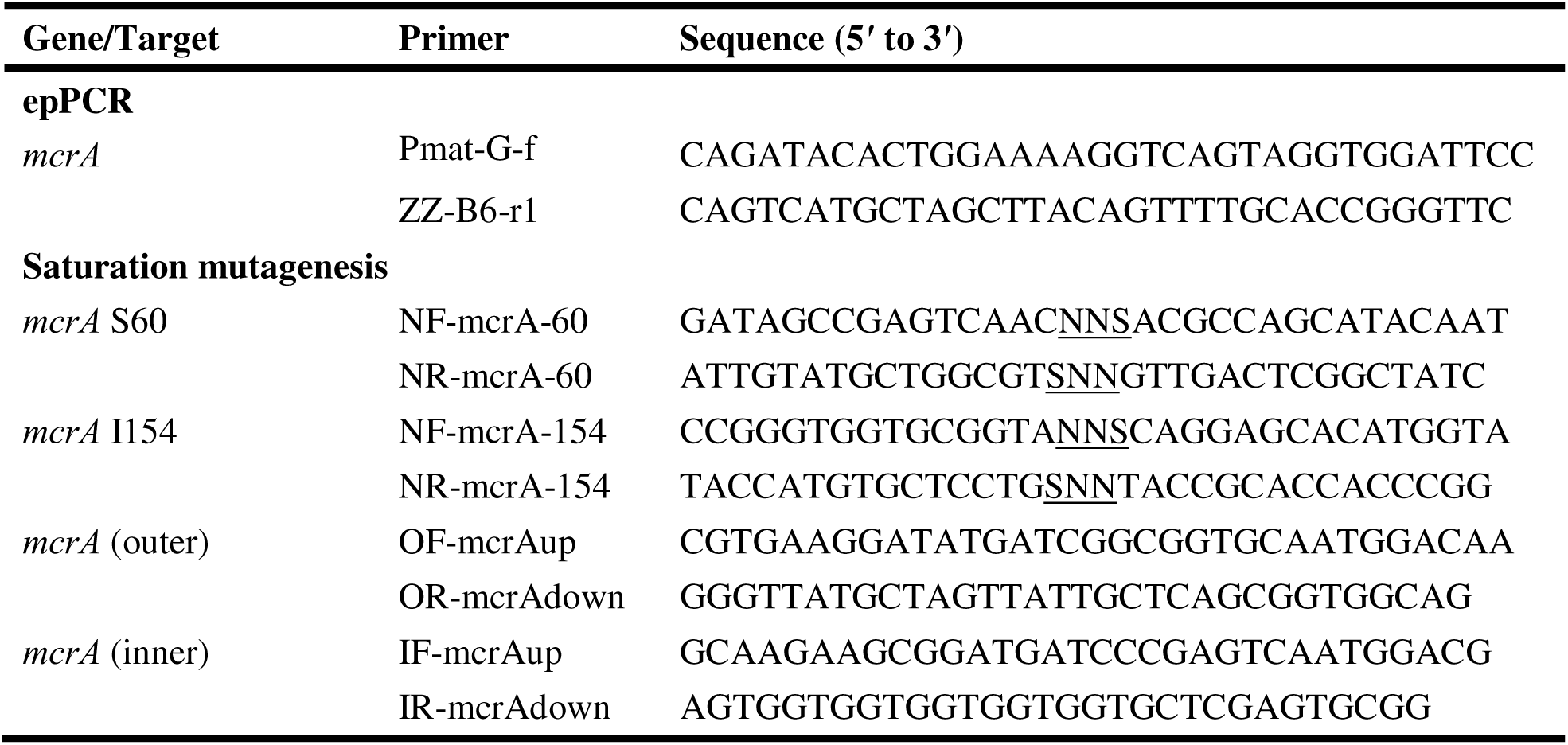
Primers used in this study. NF/NR, degenerate primers containing underlined NNS codons (N = A, T, G, or C; S = G or C); OF/OR, outer primers; IF/IR, inner primers. IF-mcrAup/IR-mcrAdown were also used to confirm transformation.

**Fig. S1.**
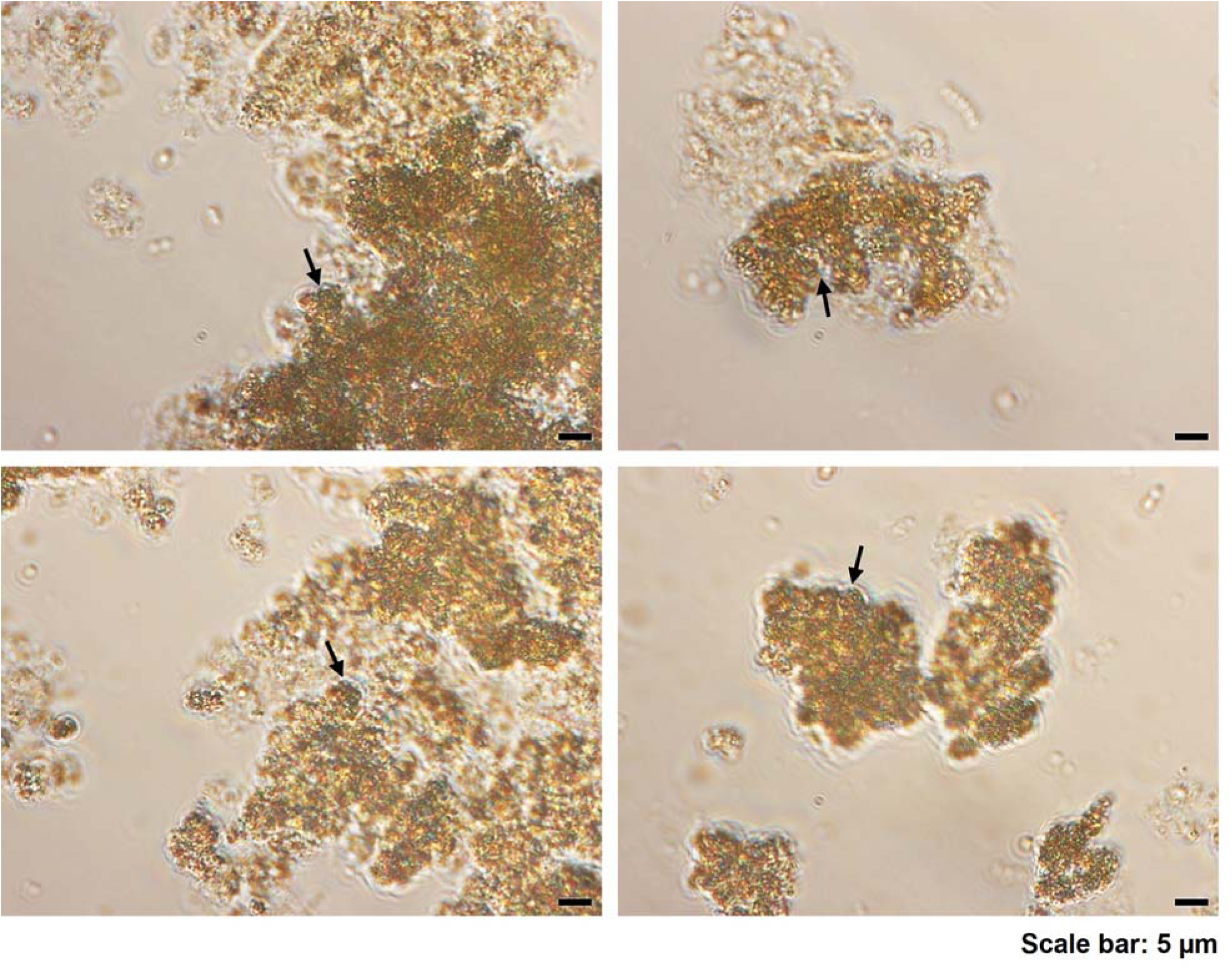
*M. acetivorans* producing Mcr_ANME-1_ from pES1-MAT*mcr3*-NcoI-mut grows as a biofilm on iron precipitates under methane culture conditions. Representative bright-field micrographs (100× magnification, Zeiss Axio Scope A1) of biofilms collected from methane-grown cultures. Cells were observed embedded within iron precipitate-associated biofilms, confirming the presence of cells within the biofilm biomass. Arrows indicate representative cells.

**Fig. S2.**
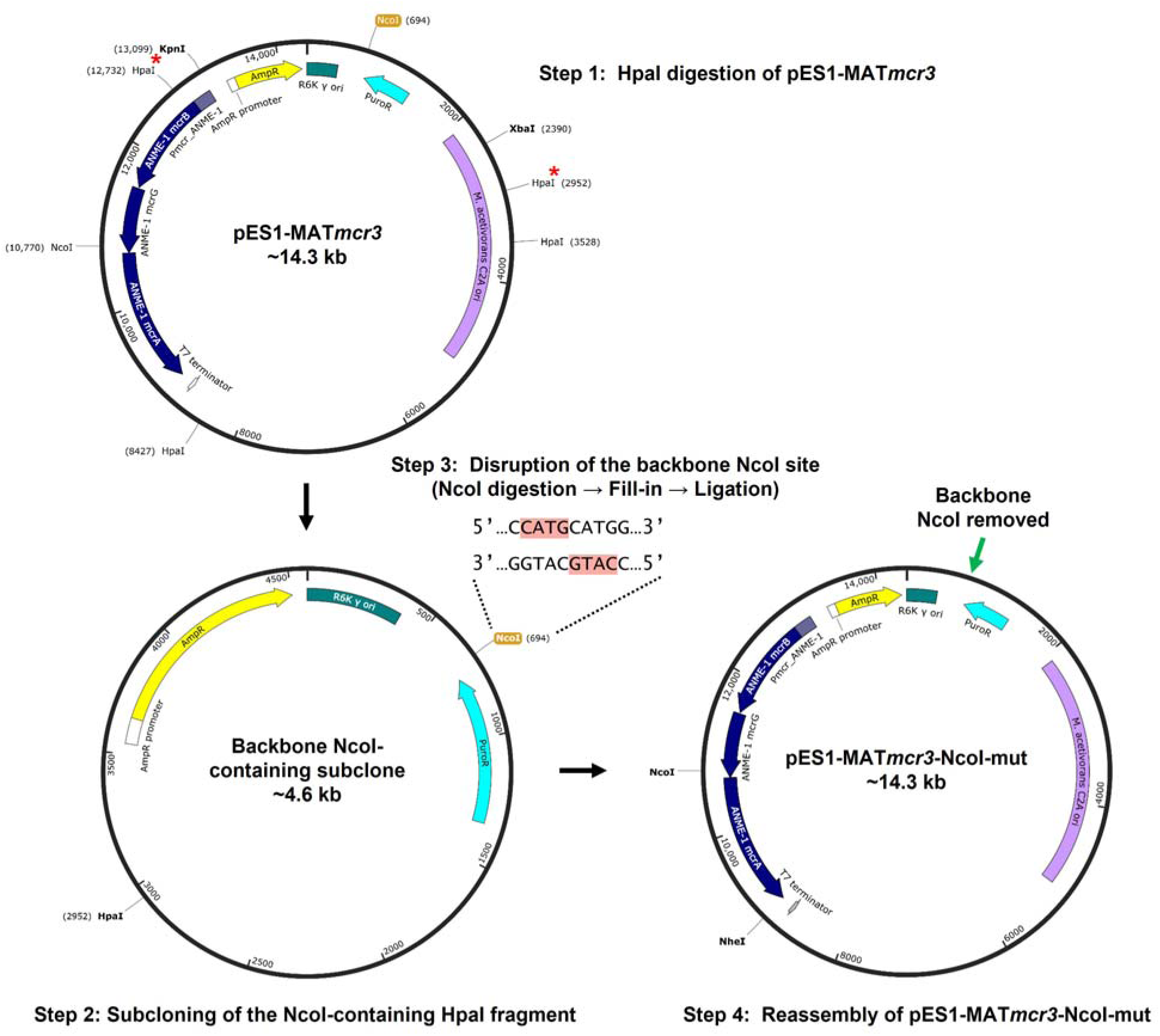
Strategy for construction of pES1-MAT*mcr3*-NcoI-mut. pES1-MAT*mcr3* contains two NcoI restriction sites, one located in the plasmid backbone (which needed to be removed; yellow) and the other located upstream of *mcrA*. To generate a unique NcoI site for cloning epPCR-mutagenized *mcrA* using NcoI and NheI, the backbone NcoI site was disrupted. Briefly, a backbone NcoI-containing HpaI fragment was subcloned (Step 2), and the NcoI site was disrupted by fill-in of the NcoI-generated 5′ overhang using T4 DNA polymerase, resulting in insertion of four nucleotides (CATG; red) (Step 3). The modified fragment was then reassembled into pES1-MAT*mcr3* using the XbaI and KpnI restriction sites to generate pES1-MAT*mcr3*-NcoI-mut (Step 4; **Fig. S3**). HpaI sites used for subcloning (Step 1) are indicated by red asterisks (*).

**Fig. S3.**
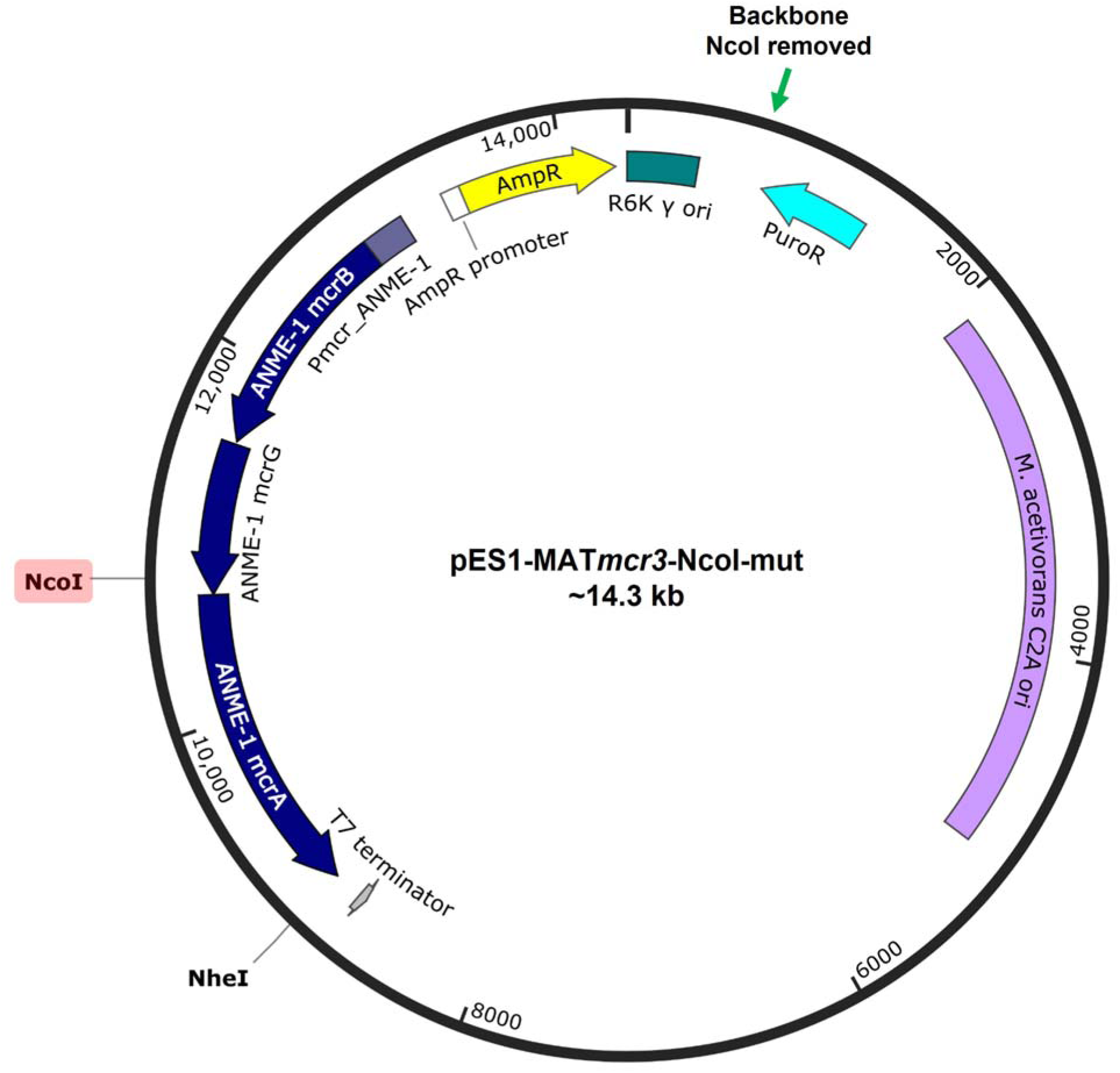
pES1-MAT*mcr3*-NcoI-mut shuttle vector used for Mcr_ANME-1_ *mcrA* epPCR library construction. In pES1-MAT*mcr3*, one NcoI restriction site is located upstream of *mcrA* (red), whereas a second NcoI site is present in the plasmid backbone (green arrow). To facilitate cloning of epPCR-mutagenized *mcrA* using the NcoI and NheI restriction sites, the backbone NcoI site was disrupted by insertion of four nucleotides (CATG) (**Fig. S2**). The resulting plasmid, pES1-MAT*mcr3*-NcoI-mut, was subsequently used for epPCR library construction. pES1 is a shuttle vector capable of replication in both *E. coli* (R6K ori) and *M. acetivorans* (C2A ori). ANME-1 *mcrBGA* is expressed under the control of its native promoter (Pmcr_ANME-1). AmpR, ampicillin resistance; PuroR, puromycin resistance.

**Fig. S4.**
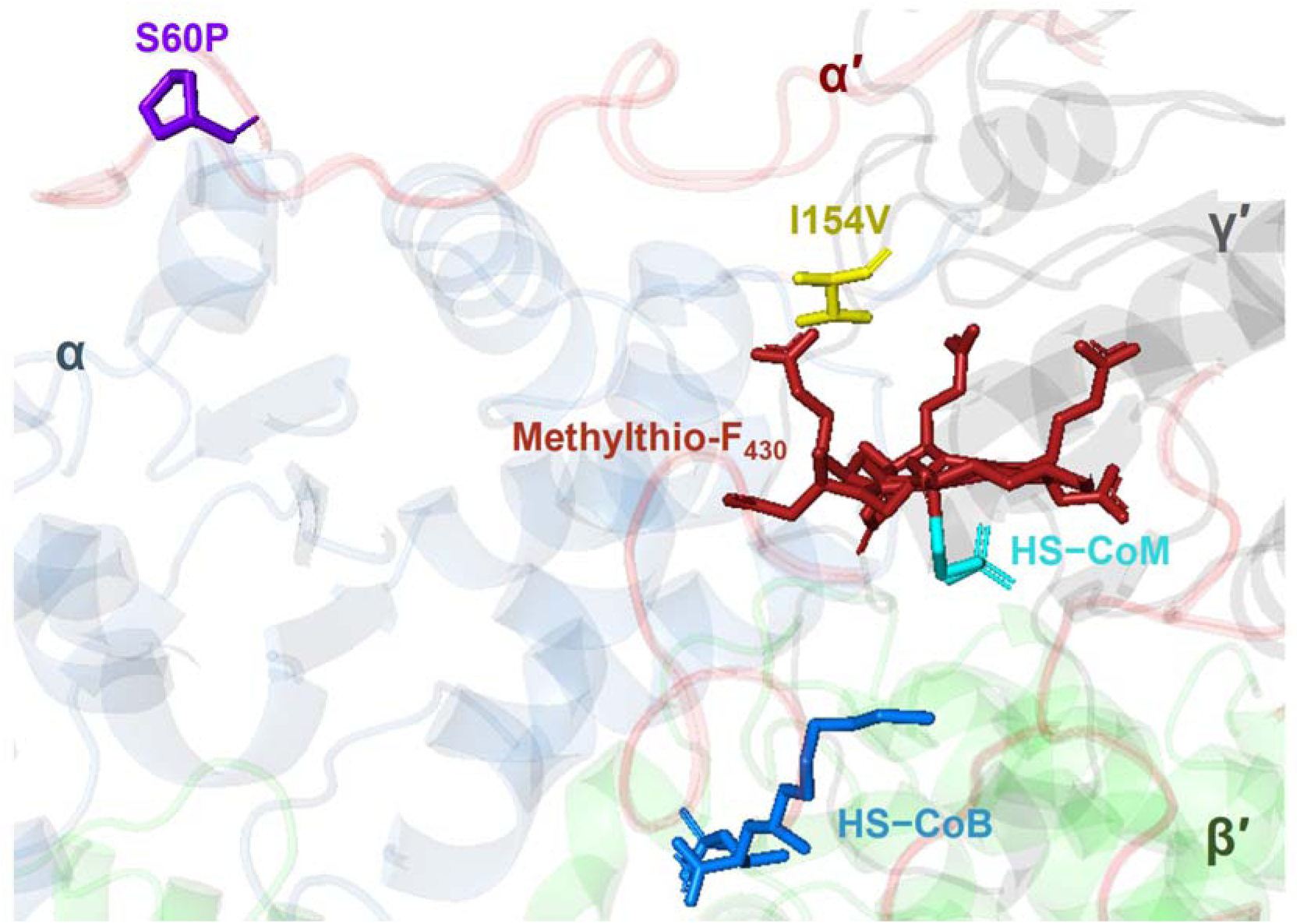
Structural context of the beneficial Mcr_ANME-1_ McrA substitutions S60P and I154V. Structural overlay of the native Mcr_ANME-1_ McrA structure and AlphaFold-predicted Mcr_ANME-1_ McrA variants carrying the beneficial substitutions S60P (purple) or I154V (yellow). The catalytic cofactor methylthio-F_430_ is shown in dark red, while HS-CoM (cyan) and HS-CoB (blue) are highlighted. The α (McrA), β (McrB), and γ (McrG) subunits are shown as cartoons in different colors for clarity. Primes (′) denote the second αβγ heterotrimer in the (αβγ)[Mcr_ANME-1_ complex. S60P is located distal to the catalytic center, whereas I154V is positioned adjacent to the methylthio-F_430_ cofactor.

**Fig. S5.**
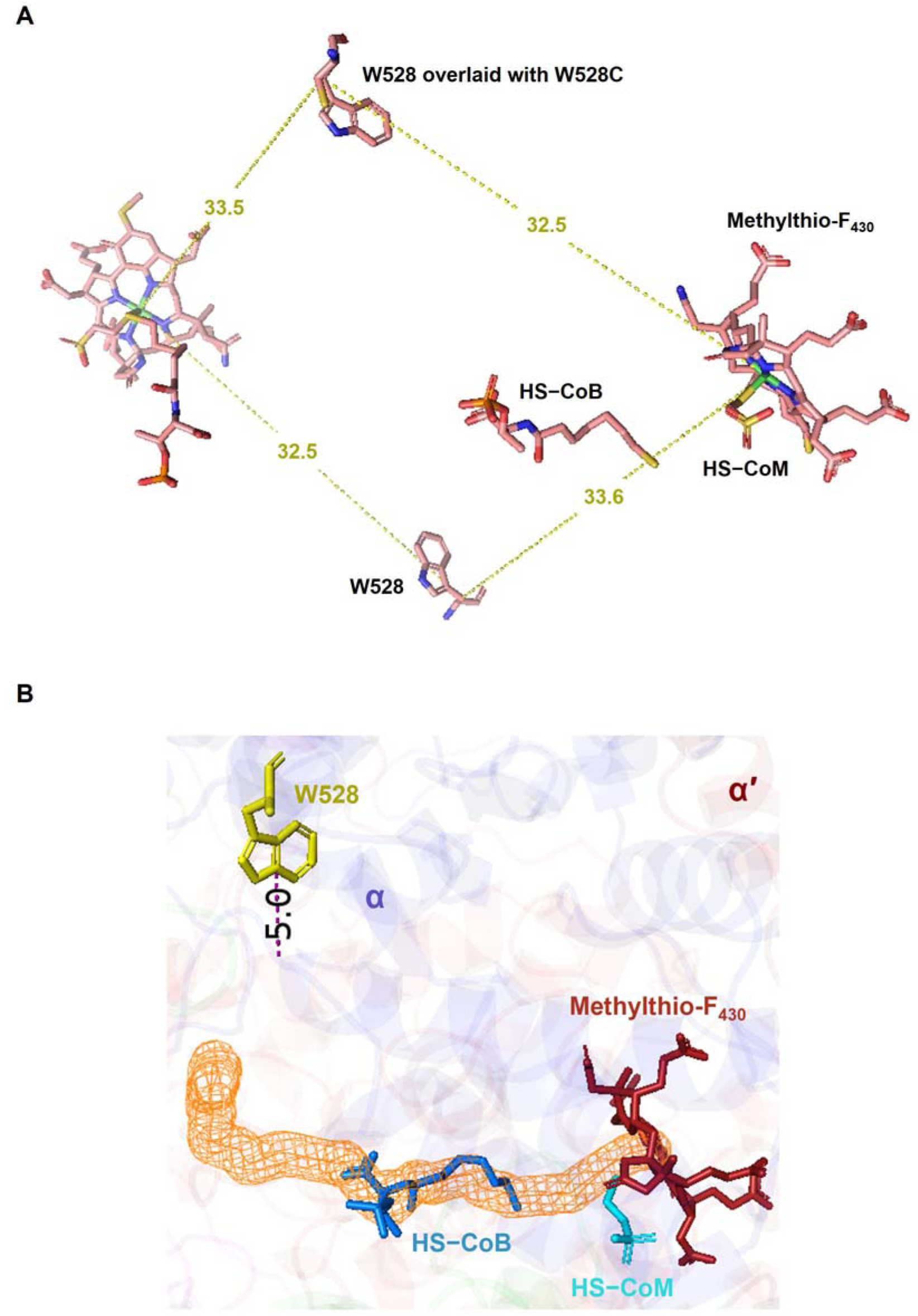
Structural context of the beneficial Mcr_ANME-1_ McrA substitution W528C. (**A**) Distances (Å) from the nickel center of the methylthio-F_430_ cofactor to the beneficial Mcr_ANME-1_ McrA substitution W528C. Carbon is shown in pink, oxygen in red, nitrogen in blue, sulfur in yellow, phosphorus in orange, and nickel in green. (**B**) Distance (Å) between Mcr_ANME-1_ McrA W528 and the nearest residue (Mcr_ANME-1_ McrA Q276) lining the tunnel (orange mesh) through which HS-CoB (blue) and CH_3_−S−CoM (cyan) exit the active site.

**Fig. S6.**
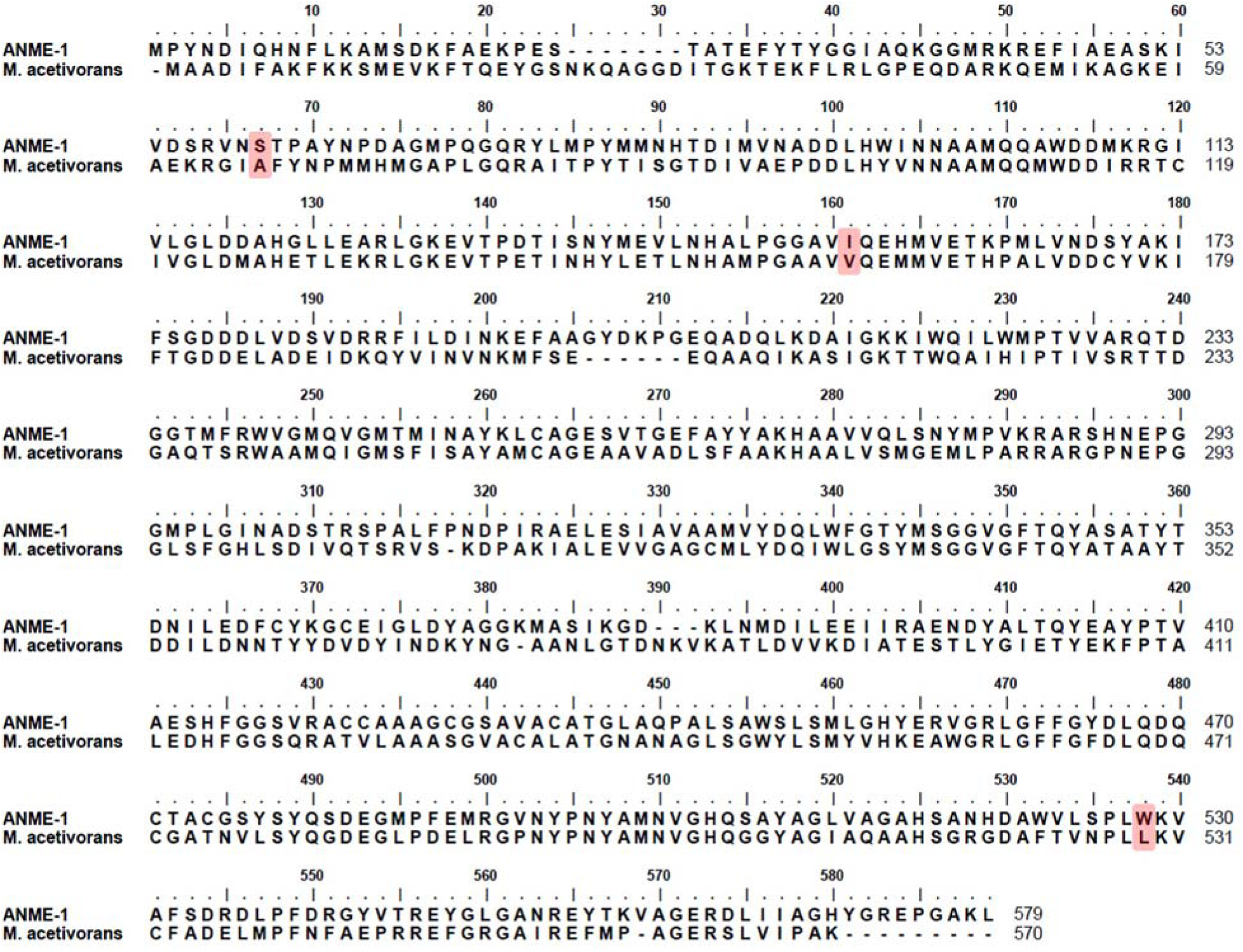
Amino acid sequence comparison of Mcr_ANME-1_ and host Mcr*_M.a._* McrA at positions corresponding to beneficial substitutions. Residues corresponding to the beneficial substitutions (S60P, I154V, and W528C) identified during laboratory evolution are highlighted in red. At positions 60, 154, and 528, Mcr_ANME-1_ contains serine (S), isoleucine (I), and tryptophan (W), whereas the corresponding residues in Mcr*_M.a._* are alanine (A), valine (V), and leucine (L), respectively. Notably, the beneficial substitution I154V restores the residue present in Mcr*_M.a._*, while S60P and W528C similarly reduce side-chain size relative to the native Mcr_ANME-1_ residues.

**Fig. S7.**
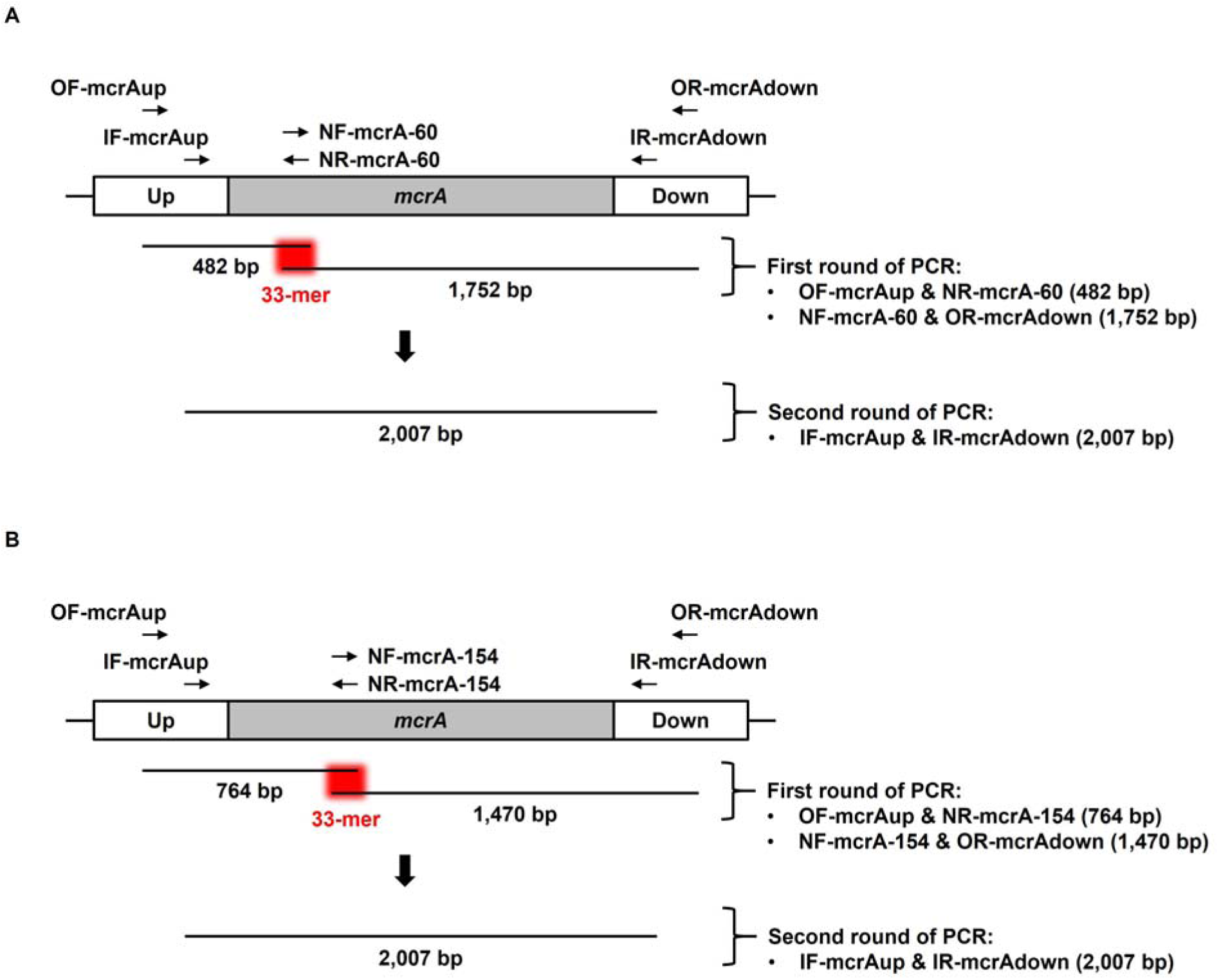
Strategy for overlap-extension PCR using nested primers for Mcr_ANME-1_ McrA S60 and I154 saturation mutagenesis. To improve PCR efficiency and facilitate library construction, a two-step PCR strategy was employed using the primers listed in **Table S3** rather than direct amplification of the entire ∼14.3 kb pES1-MAT*mcr3*-NcoI-mut plasmid. In the first round of PCR, two overlapping fragments were amplified separately: 482 bp and 1,752 bp for S60 saturation mutagenesis (**A**), or 764 bp and 1,470 bp for I154 saturation mutagenesis (**B**). The two fragments shared a 33-mer overlap containing the NNS codon used for saturation mutagenesis (red). In the second round of PCR, the overlapping fragments were assembled and amplified using the nested primers IF-mcrAup and IR-mcrAdown, generating the final 2,007 bp mutagenized product. The resulting PCR product was subsequently cloned into pES1-MAT*mcr3*-NcoI-mut using the NcoI and NheI restriction sites.

**Fig. S8.**
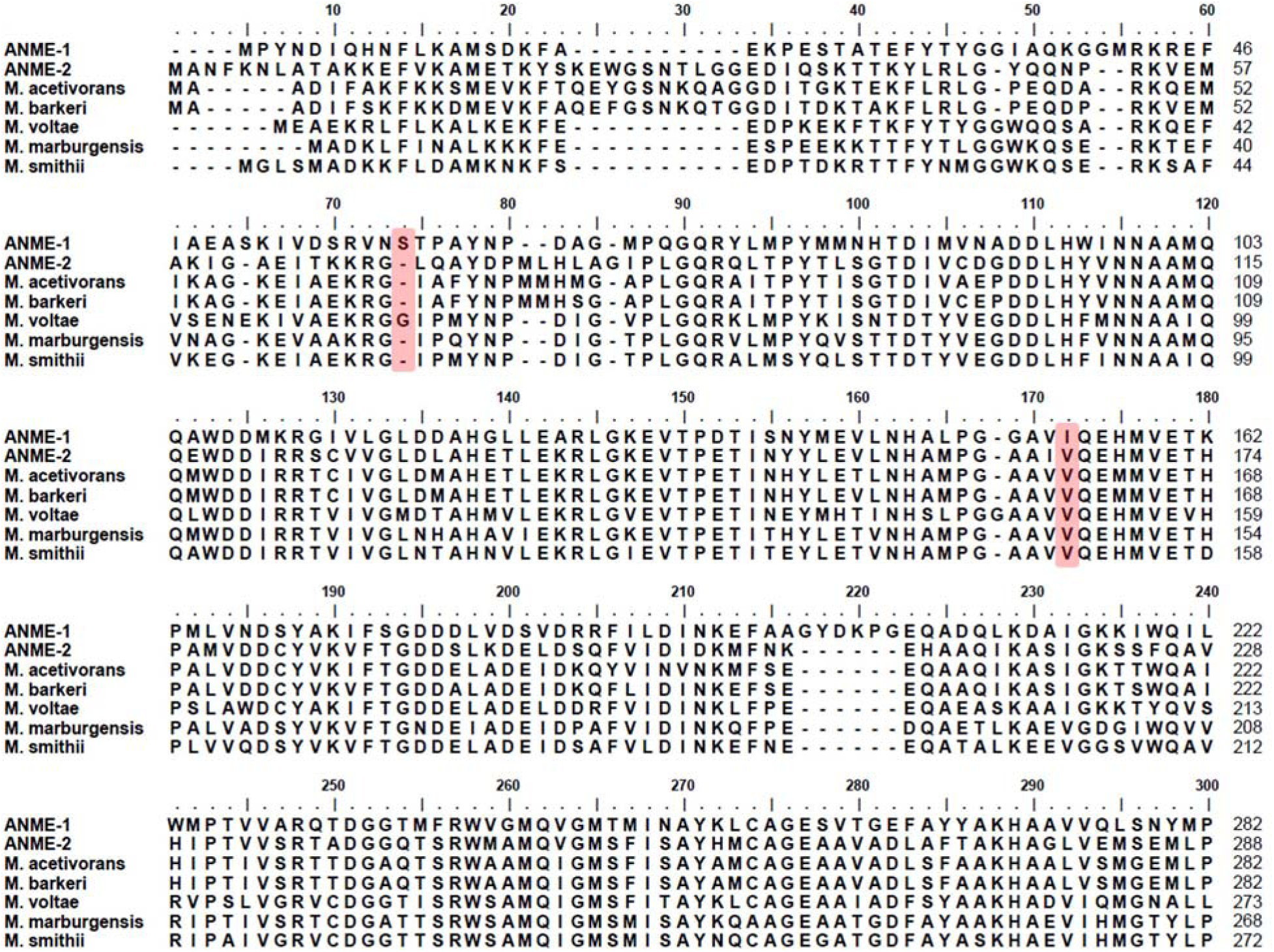
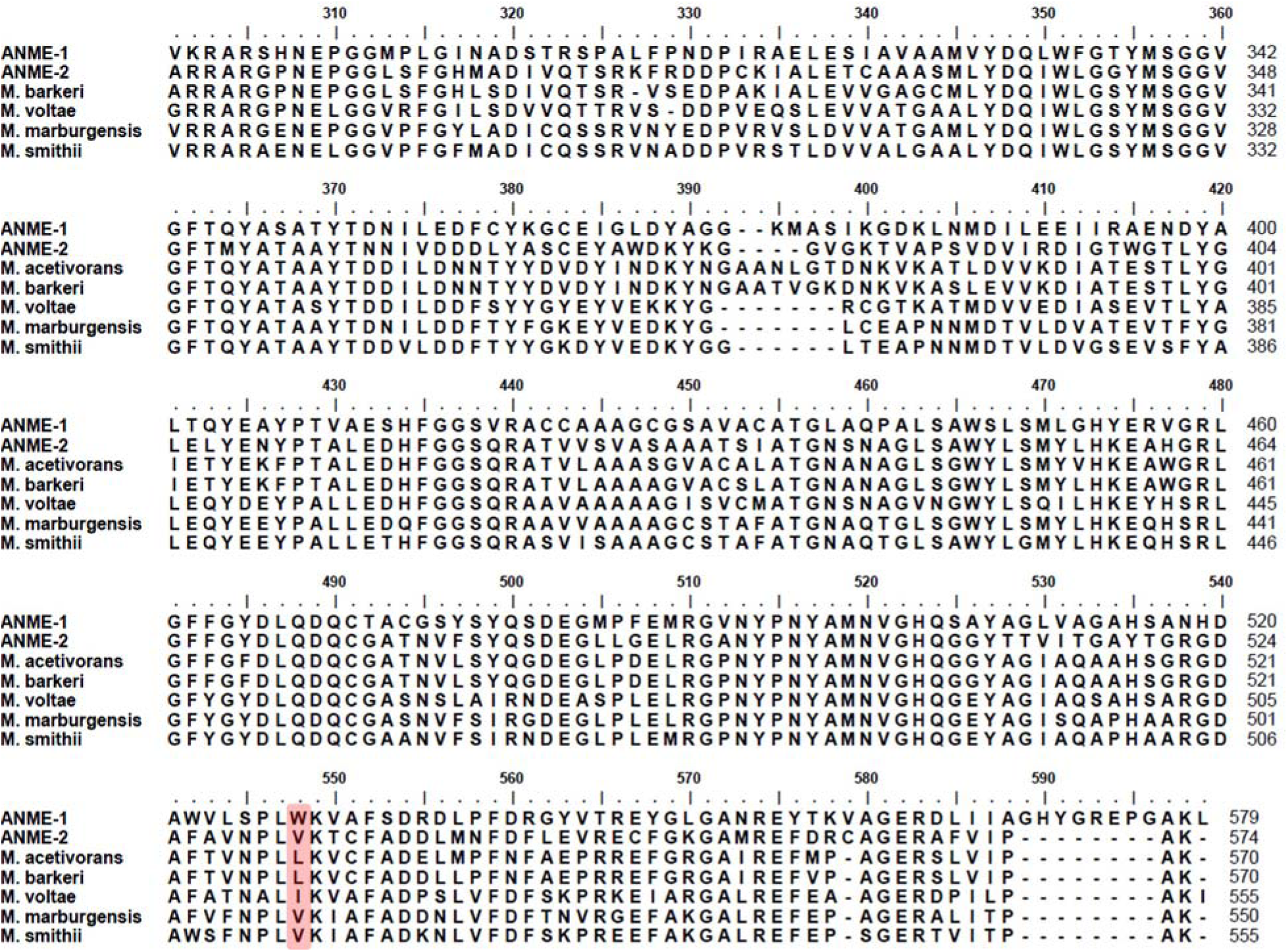
McrA amino acid sequence comparison among anaerobic methanotrophic (ANME) and methanogenic archaea. McrA amino acid sequences from ANME-1, ANME-2, *M. acetivorans*, *M. barkeri*, *M. voltae*, *M. marburgensis*, and *M. smithii* were aligned using Clustal W. Alignment positions corresponding to residues S60, I154, and W528 in Mcr_ANME-1_ McrA, identified during protein engineering, are highlighted in red.

